# DANGER ANALYSIS: RISK-AVERSE ON/OFF-TARGET ASSESSMENT FOR CRISPR EDITING WITHOUT A REFERENCE GENOME

**DOI:** 10.1101/2023.03.11.531115

**Authors:** Kazuki Nakamae, Hidemasa Bono

## Abstract

The CRISPR-Cas9 system has successfully achieved site-specific gene editing in organisms ranging from humans to bacteria. The technology efficiently generates mutants, allowing for phenotypic analysis of the on-target gene. However, some conventional studies did not investigate whether deleterious off-target effects partially affect the phenotype. Herein, we present a novel phenotypic assessment of CRISPR-mediated gene editing: Deleterious and ANticipatable Guides Evaluated by RNA-sequencing (DANGER) analysis. Using RNA-seq data, this bioinformatics pipeline can elucidate genomic on/off-target sites on mRNA-transcribed regions related to expression changes and then quantify phenotypic risk at the gene ontology (GO) term level. We demonstrated the risk-averse on/off-target assessment in RNA-seq data from gene-edited samples of human cells and zebrafish brains. Our DANGER analysis successfully detected off-target sites, and it quantitatively evaluated the potential contribution of deleterious off-targets to the transcriptome phenotypes of the edited mutants. Notably, DANGER analysis harnessed *de novo* transcriptome assembly to perform risk-averse on/off-target assessments without a reference genome. Thus, our resources would help assess genome editing in non-model organisms, individual human genomes, and atypical genomes from diseases and viruses. In conclusion, DANGER analysis facilitates the safer design of genome editing in all organisms with a transcriptome.

## INTRODUCTION

The CRISPR-Cas9 system was initially adapted as a bacterial immune system(^1,2^). Over the past decade, this system has been developed as a programmable nuclease that enables site-specific modification of the genomes of various organisms, including humans)^3–5^), insects)^6,7^), microalgae(^8,9^), and bacteria(^10^). Engineered CRISPR-Cas9 undertakes genomic modification using two components: RNA-guided Cas9 nuclease and single-guide RNA (sgRNA)(^11,12^). The Cas9-sgRNA complex generates indels near the target site (on-target site), where the 19–20 bases of the 5´ ends of sgRNA (protospacer) and the protospacer adjacent motif (PAM) of Cas9 protein bind(^11–14^). Recently, many CRISPR-Cas9 applications, such as Cas9 nickase (Cas9n)(^12^), dead Cas9 (dCas9)(^15^), base editors(^16,17^), and prime editors(^18^), have been developed. Furthermore, CRISPR-Cas9-mediated genome editing was found to be efficient, with the editing efficiency exceeding 50% over time(^19^). Thus, CRISPR technology has dramatically facilitated a reverse genetics approach involving phenotypic analysis using CRISPR-Cas9-based mutants of a user-targeted gene(^20–23^).

However, genome editing using CRISPR technology presents two challenges that have not been addressed in previous studies. First, phenotypic effects caused by unexpected CRISPR dynamics are not quantitively monitored. CRISPR-Cas9 is well known for unexpected sequence editing (off-target site) with mismatches when compared to protospacers and PAM^5^. Off-target gene editing results in incorrectly edited mRNA, unexpected phenotypes, and decreased expression of unrelated genes. Some reports predicted and detected off-target editing using genomic PCR and DNA sequencing analysis(^14,24–26^), but most studies have not assessed the phenotypic effect of the detected off-targets. Second, CRISPR technology generally depends on basic genomic data, including the reference genome. CRISPR technology has potential applications in organisms with incompletely characterized genomes. However, the design of site-specific sgRNAs requires the factual genomic sequence of materials to be treated with CRISPR technology. This hindrance also emerges in the human genome, particularly in the genomes of patients and cancer genomes. These genomes are assumed to be completely distinct from the reference genome(^27,28^). The off-target is always “unexpected.” Thus, we need a method to observe factual genomic sequences and reduce potential off-target effects.

We devised a method to overcome the two challenges above: phenotypic risk and dependence on a reference genome. Phenotypic risk can be assessed by phenotype analysis using gene ontology (GO) annotation(^29,30^). GO has been widely used for several decades(^20,23,31–34^). Recently, many RNA sequencing (RNA-seq) data and mapped genes have been annotated with GO terms to characterize the transcriptome phenotype under a specific condition of the organism. This process is known as enrichment analysis(^35^). We expected that we could quantitatively assess the phenotypic risk of off-target genes if each off-target gene with decreased expression was annotated with GO terms. Moreover, *de novo* transcriptome assembly technology can address the dependency problem of reference genomes. The *de novo* transcriptome assembly can generate transcriptome sequences without the reference genome using RNA-seq data(^36–40^). We identified factual genomic sequences in mRNA-transcribed regions using *de novo* transcriptome assembly from gene-edited organisms and cells.

In this study, we combined *de novo* transcriptome assembly and GO annotation analysis in CRISPR editing to establish a DNA on/off-target assessment, including phenotype risk analysis without a reference genome. We named it **D**eleterious and **AN**ticipatable **G**uides **E**valuated by **R**NA-sequencing (**DANGER**) analysis (Figure 1). This bioinformatics pipeline can elucidate genomic on/off-target sites based on *de novo* transcriptome assembly using RNA-seq data. Then, it identifies the *deleterious off-targets*, defined as off-targets on the mRNA-transcribed regions that represent the downregulation of expression in edited samples compared to wild-type (WT) ones. Furthermore, our pipeline can quantify phenotypic risk at the GO term level by calculating a newly defined indicator of phenotypic risk by the deleterious off-targets, named the D-index.

**Figure 1.**
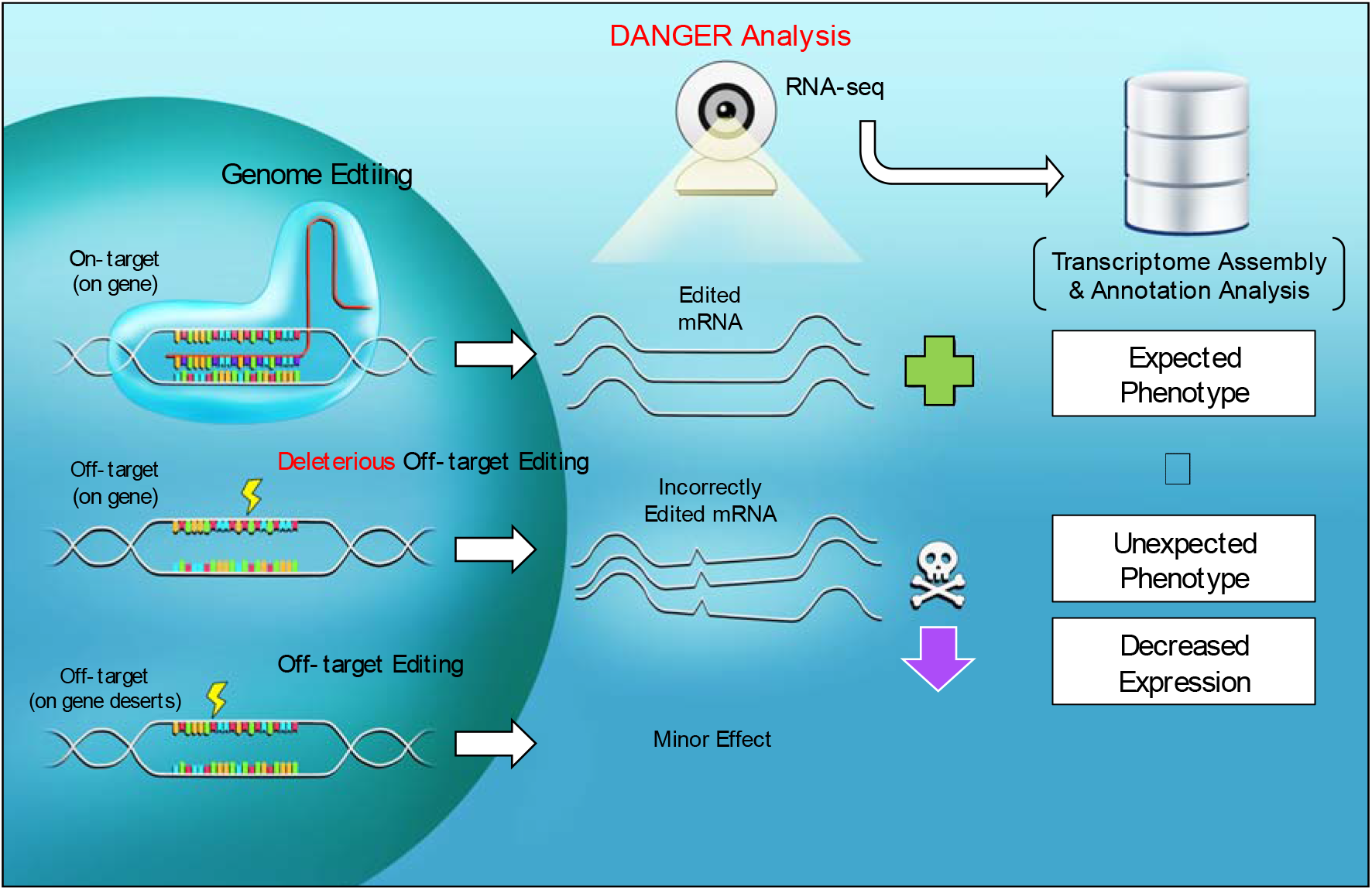
Scheme of CRISPR-Cas9 targeting, deleterious off-target editing, and DANGER analysis.

## MATERIAL AND METHODS

### Implementation of DANGER Analysis

Our pipeline of DANGER analysis is composed of several processes: "Quality control & Adapter trimming," "rRNA Removal," " *de novo* transcriptome assembly," " Removal of redundancy," "Detection of on-target and potential off-target sites," "Expression quantification," "Search for deleterious off-target sites," "Identification of ORFs and Genes," "GO analysis," and "Validation for phenotypic risk" (Figure 2A).

**Figure 2.**
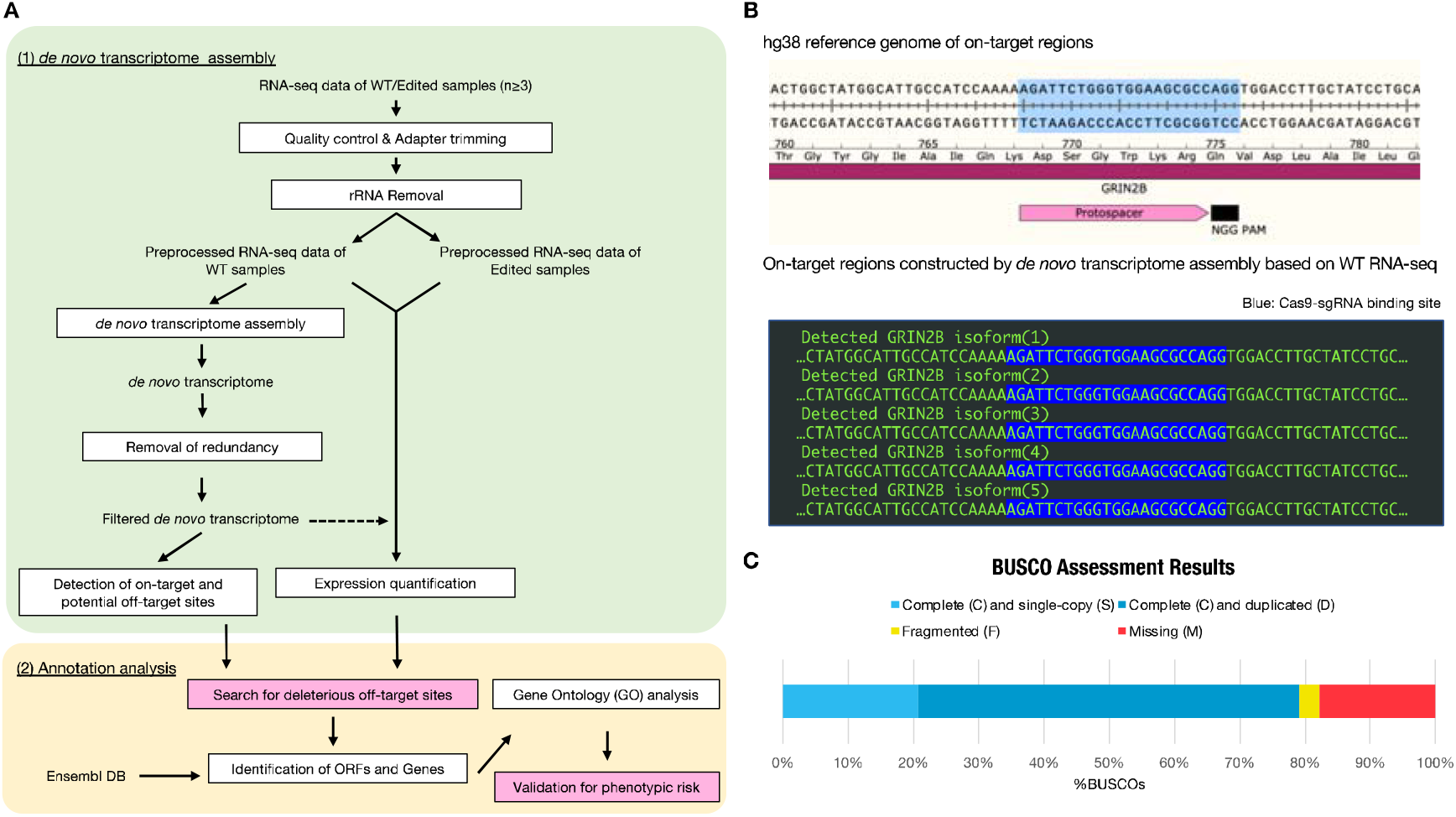
Overview of DANGER analysis and on-target region constructed by *de novo* transcriptome assembly. A. Bioinformatic workflow of DANGER analysis. Our analysis requires RNA-seq data derived from WT and Edited (each n≥3). DANGER analysis has two steps in the workflow: (1) *de novo* transcriptome assembly (light green background color) and (2) annotation analysis (light yellow background color). The *de novo* transcriptome assembly step is processed with Trinity and preprocessing tools such as cutadapt and bbduk.sh. Crisflash performs the search of on/off-target sequences. The RSEM quantifies gene expression in Edited RNA-seq samples in comparison to the WT *de novo* transcriptome (dot allow). The step of annotation analysis was involved processing with TransDedoder, ggsearch, org.XX.eg.db (e.g., org.Hs.eg.db in the transcriptome related to humans), and topGO. We implemented specific modules, colored in pink, for considering the phenotypic effect of deleterious off-targets. B. Comparison between the hg38 reference genome and transcript sequence constructed by *de novo* assembly of RNA-seq samples derived from WT iPSC-derived cortical neurons on the GRIN2B on-target region. The on-target region of the hg38 reference genome is illustrated with annotations of the *GRIN2B* CDS, the protospacer, and the NGG PAM sequence of SpCas9. The detected GRIN2B isoforms (1–5) are lined up in green in the black box. The Cas9-sgRNA binding sites are highlighted in blue. C. Genome completeness of *de novo* transcriptome assembly RNA-seq data derived from WT iPSC-derived cortical neurons was assessed using conserved mammal BUSCO genes (mammalia_odb10). The result was 79.1% of “complete,” 20.7% of “single-copy,” 58.4% of “duplicated,” 3.2% of “fragmented,” and 17.7% of “missing” (n = 9226).

DANGER analysis examines paired-end RNA-seq data derived from wild-type (WT) and edited samples using the processes depicted in Figure 1. The pipeline generates a *de novo* transcriptome assembly, an expression profile of transcripts belonging to on/off-target sites, and an estimation of the phenotypic risk for off-target sites. The script has been uploaded to our GitHub repository (https://github.com/KazukiNakamae/DANGER_analysis). The analyses were performed on Docker with Ubuntu v. 22.04.1, LTS, and 235 GB of memory. The scripts for this processing pipeline were released as a Docker image (https://hub.docker.com/r/kazukinakamae/dangeranalysis), enabling operation on various operating systems beyond Linux using this Docker image. Each process is explained in detail below.

#### Quality Control & Adapter Trimming

Quality control and adapter trimming were performed using Cutadapt v. 1.18(^41^), which also removed low-quality reads. The adapter sequences used were "AGATCGGAAGAG."

#### Ribosomal RNA (rRNA) Removal

The residual rRNA reads were filtered using bbduk v. 38.18 (https://sourceforge.net/projects/bbmap/). Each sample was filtered twice. In the first and second filters, we used SSU and LSU rRNA datasets from SILVA v. 119.1 (https://www.arb-silva.de). The dataset was downloaded from the CRISPRroots (https://rth.dk/resources/crispr/crisprroots/).

#### De novo Transcriptome Assembly

The *de novo* transcriptome assembly was performed using Trinity v. 2.12.0(^38^). The merged read files were composed of RNA-seq data derived from WT samples. Transcriptome completeness was assessed using BUSCO v. 5.2.2_cv1(^42^). The BUSCO evaluated the competence of assembly using estimation of similarity to gene database (BUSCO genes) and classified hit sequence into “complete” (including “single-copy” and “duplicated”), “fragmented,” and “missing.” The databases used for conserved mammalian BUSCO genes were "mammalia_odb10" and "actinopterygii_odb10" for human and zebrafish assemblies, respectively.

#### Removal of redundancy

The expression of RNA-seq data derived from WT samples was quantified in advance to remove transcripts with low expression levels. Quantification was performed with align_and_estimate_abundance.pl of Trinity v. 2.12.0. The removal of transcripts was performed with filter_low_expr_transcripts.pl of Trinity v. 2.12.0.

#### Detection of on-target and potential off-target sites

Detection of on/off-target sites in *de novo* transcriptome assembly was performed with Crisflash v.1.2.0(^43^). Our off-target detection focused on up to 8 or up to 11nt mismatches and NGG or NRR PAM because the previous off-target reports indicated that off-target sites with ≥5 nt mismatches and NGG/NAG/NGA/NAA PAM exist^24,44^, although not with a high frequency. Specifically, we searched for potential off-target sites by executing the following command:

“crisflash -g Result_denovo_transcriptome_ { *de novo* transcriptome assembly generated from WT RNA-seq data} -s {sequence of protospacer and PAM} -o {output file of Crisflash} -m {the maximum number of mismaches users consider} -p {PAM} -t {the number of thread} -C.“

The output file of Crisflash has on-/off-target locations, off-target sequences, and mismatch numbers included in the tab-delimited format (Cas-OFFinder format). The on-target locations were extracted using perfect matching with the expected genomic sequence of Cas9-sgRNA binding. The potential off-target sites were classified by the mismatch number in each text file.

The detection of on/off-target sites uses one *de novo* transcriptome assembly generated by merging all replicates of RNA-seq data from WT samples to ensure the best quality of assembly. Thus, we did not obtain replicates of numerical data related to off-target sites or conduct statistical evaluations such as p-values using the replicated data. The analysis design is due to concerns that utilizing *de novo* transcriptome assembly generated from individual replicates might diminish the accuracy of the analysis.

#### Expression quantification

Using filtered *de novo* transcriptome assembly, we quantified the expression of RNA-seq data derived from each sample. Quantification was performed using align_and_estimate_abundance.pl of Trinity v. 2.12.0. The transcripts per million (TPM) dataset was constructed from “RSEM.isoforms.results” of each output directory and was saved as a single CSV file.

The ratio of TPM values on a transcript was calculated with the following formula:

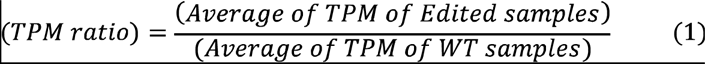

A transcript with a TPM ratio of less than the user-defined value (t) was determined as downregulated TPM (dTPM). Furthermore, in place of TPM, it is compatible with profiling based on Differentially Expressed Genes (DEG). The DEG analysis was performed to compare different analysis methods. The read count data were extracted from the "RSEM.isoforms.results" of each output directory in the section "Implementation of DANGER analysis: Expression quantification.” The raw count data were normalized by the Tag Count Comparison (TCC) R package(^45^) with the parameter "norm.method="tmm," test.method="edger," iteration=30, FDR=0.1, floorPDEG=0.05" to detect DEG between RNA-seq data derived from WT and Edited samples. The MA plot was constructed using an in-house R script. If a DEG transcript had a negative log-ratio of normalized counts (M-value) and its p-value fell below the user-defined value (α), it was determined to be downregulated DE (dDE). It was saved as a single CSV file.

#### Search for deleterious off-target sites

Our pipeline defined an off-target site where the transcript was annotated with dTPM or dDE as a deleterious off-target site, which could be also paraphrased as “actual off-target site.” We counted the number of deleterious off-target sites using an in-house Python script, which required the off-target site profile and the TPM ratio or DEG described above.

#### Identification of ORFs and Genes

Our pipeline identifies open reading frames (ORFs) in the filtered *de novo* transcript sequences and predicts the corresponding amino acid sequences. If these predicted sequences exhibit significant homology with protein sequences in a database, we assign them as genes. The open reading frames (ORFs) and genes of the filtered *de novo* transcript were estimated with TransDecoder v. 5.5.0 and ggsearch v. 36.3.8g using a protein database. The process was based on the Systematic Analysis for Quantification of Everything (SAQE) pipeline (https://github.com/bonohu/SAQE)(^33^). In particular, "11TransDecoder.sh", "12GetRefProts.sh", "15ggsearch.sh", "15parseggsearch.sh", and "15mkannotbl.pl" were used with the supplemental script "00_prepare_faa_4Fanflow. sh" in the GitHub repository (https://github.com/RyoNozu/Sequence_editor). The referred protein databases were the Ensembl databases of all translations resulting from Ensembl genes in humans and zebrafish. The DANGER analysis database can be manually customized with the organism from which the analyzed RNA-seq data was extracted.

The above identification was based on one *de novo* transcriptome assembly generated by merging all replicates of RNA-seq data from WT samples to ensure the best quality of assembly. Thus, we did not obtain replicates of numerical data related to the identifications or conduct statistical evaluations such as p-values using the replicated data. The analysis design is due to concerns that utilizing *de novo* transcriptome assembly generated from individual replicates might diminish the accuracy of the analysis.

#### GO enrichment analysis

GO annotations of gene ontologies were performed using an in-house R script using the org.Hs.eg.db and org.Dr.eg.db R packages for humans and zebrafish, respectively. Enrichment analysis followed by GO annotations was performed using the topGO R package against genes whose off-target sites were determined to be deleterious off-target sites. Enrichment analyses were performed per off-target mismatch number. Finally, the enrichment tables were merged into a single table, named the DANGER table, with the mismatch number annotated.

#### Validation for phenotypic risk

We defined the following value (D-index) to evaluate the phenotypic risk posed by deleterious off-target effects per GO term.

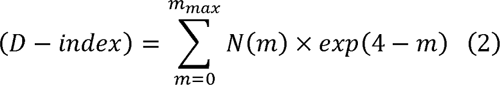

where m indicates the number of mismatches. The N(m) represents the total number of genes, which have m bases mismatches, included in a specific GO term. The m _max_ is the maximum number of mismatches that a user considers. The D-index considers both the phenotypic risk and the frequency of off-target effects. Based on previous reports, we used the exponential function to express the frequency of off-targets because the frequency of off-targets tends to decrease exponentially as the mismatch number of off-target sites increases(^24,46^). Based on a GUIDE-seq study^24^, most reported off-targets possess mismatches of four or fewer nucleotides (Supplementary Table S1). Therefore, the exponent value is represented as a decreasing function by subtracting the number of mismatches from four, and by minimizing the impact of mismatches with five or more nucleotides, we have enabled the probabilistic risk assessment at an appropriate level. Calculations were performed using an in-house Python script.

### Validation of D-index

We established a validation methodology of statistical significance for the D-index, determined as described above, for each set of GO terms using a permutation test. The permutation test involved random shuffling of the expression profile and off-target profile based on a given seed value, and the D-index was computed based on these randomly shuffled profile data. We will refer to this D-index as a pseudo-D-index. After repeating this process of creating pseudo-D-indexes 100 times, we made a distribution of pseudo-D-indices for each set of GO terms. A D-index outside the (1 – L) ×100 % confidence interval of this distribution and higher than the mean value was defined as a “significant D-index.” Additionally, we implemented a script to measure the false positive rate of the permutation test. In false-positive detection, ten additional shuffled data sets were generated using seed values different from the ones used to create the shuffled expression and mismatch profiles, and the newly calculated pseudo-D indices from these data were calculated. If they met the criteria for a significant D-index in the above distribution, the pseudo-D-indices were defined as “significant pseudo-D-index” and counted. The ratio of the significant pseudo-D-indices to the total number of new pseudo-D-indices was defined as the false positive rate. Multiplying this false-positive rate by the total number of original D-indices allows us to estimate the number of expected false D-indices, and subtracting this from the total number of original D-indices enables us to count the number of expected true D-indices.

The analyses were performed on Docker with Ubuntu v. 22.04.1, LTS, and 235 GB of memory. The scripts for this processing pipeline were released as a Docker image (https://hub.docker.com/r/kazukinakamae/dangertest), enabling operation on various operating systems beyond Linux by using this Docker image.

### Datasets

Two RNA-seq datasets were analyzed to evaluate our pipeline. First, we collected paired-end RNA-seq datasets, which had a total of >100 M reads for all WT samples and an average length of >100 nt, to ensure good quality, which indicated a percentage of complete benchmarking universal single-copy orthologs (BUSCO) genes of >70%, of transcriptome completeness after *de novo* transcriptome assembly. The datasets were downloaded from the Sequence Read Archive (SRA) (Table 1).

**Table 1.**
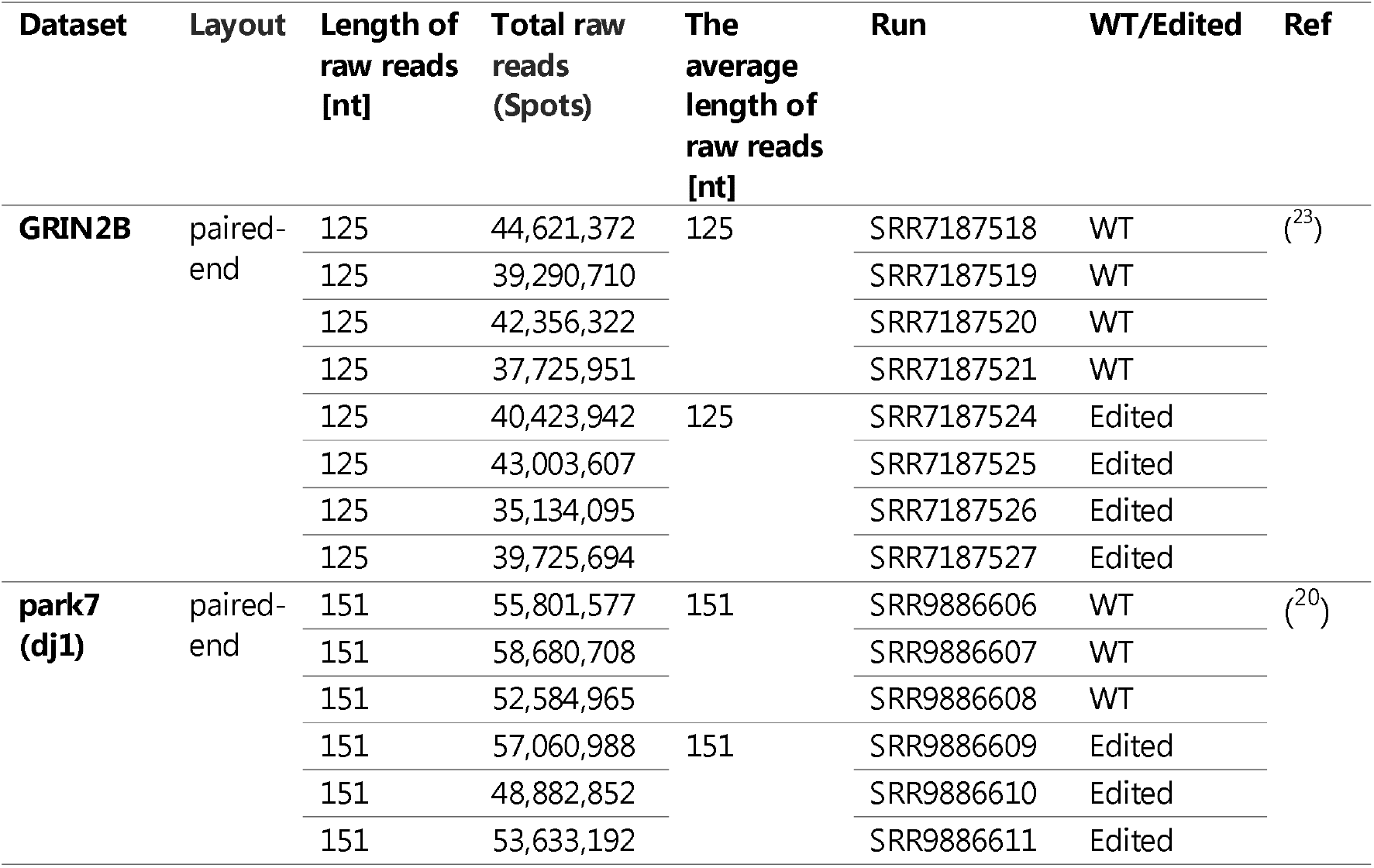
RNA-seq datasets evaluated for testing the DANGER analysis pipeline.

#### GRIN2B

The dataset was extracted from human iPSC-derived cortical neurons with or without indels generated by paired Cas9 nickase (Cas9n)-single-guide RNA (sgRNA) (GRIN2B-FW and GRIN2B-REV sgRNAs) on the GRIN2B locus(^23^). Previous studies have established clones with indels resulting in loss-of-function (LOF) and reduced dosage (RD). However, we focused only on LOF samples that had been edited and analyzed using our pipeline. Gorodkin and Seemann have previously reported that off-target sites affect the expression profile of LOF samples using reference-based RNA-seq analysis. Our study used the GRIN2B dataset to benchmark *de novo* transcriptome assembly based and reference-based RNA-seq analyses(^44^). Moreover, we profiled the on/off-target assessment of GRIN2B-REV sgRNA.

#### Park7

The dataset was extracted from the zebrafish brain with or without biallelic indels generated by a single Cas9-sgRNA at the park7 locus (which encodes DJ-1)(^20^). The analyzed F2 mutants were generated from a cross between two heterozygous F1 mutants. We used the dataset as an *in vivo* example of the DANGER analysis pipeline in a simple CRISPR-Cas9-mediated knock-out experiment.

### Statistical analysis

Plots were made using Microsoft Office and housemade Python scripts. The Exact Fisher’s test was performed for the p-value was calculated accordingly using *fisher_exact()* of the *scipy* package in Python. The 2-tailed Welch’s t-test was performed for the p-value was calculated accordingly using *ttest_ind(equal_var=False)* of the *scipy* package in Python. We used G power software for the statistical power (1-ß) calculation. The Venn diagrams were generated using web software (“Calculate and draw custom Venn diagrams”: https://bioinformatics.psb.ugent.be/webtools/Venn/).

## RESULTS & DISCUSSION

### Assessment of CRISPR-Cas9 off-targets using DANGER Analysis for RNA-seq Data from *in vitro* Differentiated Human iPSC

We investigated whether our DANGER analysis pipelines could detect deleterious off-target sites without information on the reference genome. First, we applied DANGER analysis to the GRIN2B dataset, which was extracted from human iPSC-derived cortical neurons with or without in-frame deletions at the GRIN2B locus(^23^). The obtained *de novo* transcriptome assembly contained five isoforms in which on-target sequences of sgRNA (GRIN2B-REV) (Figure 2B) were located. The assembly comprised 342,910 contigs and exhibited BUSCO transcriptome completeness of 79.1% (Figure 2C). Previous studies on *de novo* transcriptome assembly using Trinity reported 64.7%, 77.1%, 80%, and 87% complete BUSCO genes in higher animals such as *Homo sapiens* (^36^), *Castor fiber* L.(^37^), *Mirounga angustirostris* (^39^), and *Dromiciops gliroides* (^40^), respectively. We successfully obtained a *de novo* transcriptome with standard quality. Consequently, the pipeline performed an exhaustive search using Crisflash, yielding 33,878 potential off-target sites with up to 8 bases mismatches (MM) and NGG PAM, using transcriptome assembly.

Next, we quantified the transcriptome-wide expression of each of the four RNA-seq samples from the WT and Edited GRIN2B loci. Our pipeline examines whether the potential off-target site, which is output of Crisflash, is in the transcript and whether it also reduces the expression value. The pipeline considers those potential off-targets that have confirmed down-expression as deleterious off-targets, in other words, actual off-targets. The reduction of expression may occur because nonsense-mediated mRNA decay (NMD)^47^ destroys incomplete transcript sequences resulting from off-targeting, or alignment software fails to map the incomplete transcript sequences to the untreated transcript sequence^48^. The DANGER analysis screens transcripts with lower expression levels in edited samples compared to WT samples. In general, there are several criteria for estimating expression using RNA-seq. We implemented two criteria, "downregulated different expression (dDE)" and “dTPM," for the detection of downregulated transcripts (Figure 3A; two callouts). Our dDE criterion uses DEG analysis with TCC normalization (see Materials and Methods). In the case of DEs in a transcript with a negative M-value and a p-value that was less than the threshold value (α) in the MA plot (Figure 3A; right callout), we defined the transcript as "dDE." On the other hand, the “TPM" criterion uses the normalized value, named TPM, as first defined by (^49^) and calculates the ratio of TPM between WT and the edited samples. When the ratio of a transcript is less than the threshold value (t), we defined the transcript as "dTPM" (Figure 3A; left callout). We confirmed the number of transcripts with dDE (α = 0.001) or dTPM (t = 0.4) annotation that contained off-target sites (Figure 3A; Venn diagram). There were 730 transcripts with dDE off-targets. A total of 12,747 transcripts with dTPM off-targets were detected, which was approximately 17-fold more than those with dDE off-targets. Our DANGER analysis aims to serve as a screening tool that emphasizes maximizing the estimation of potential risks by capturing as many sites suspected of phenomena as knockout or knockdown of off-target genes. From this perspective, the dTPM approach can estimate the deleterious off-target effects to the greatest extent, more so than the dDE approach. Furthermore, we focused on the off-targets detected in common by dTPM and dDE. We observed their mismatch count (Supplementary Figure S1A). As a result, we confirmed that the off-targets seen in common were distributed at an even ratio for each mismatch count compared to the off-targets detected only by dTPM (Supplementary Figure S1B). Given that the number of mismatches affects the likelihood of off-target occurrences, there is no correlation between the common off-targets and their occurrence rate. While there are differences in the number and types of transcripts detected by dTPM and dDE, no evidence suggests that these differences result from a flaw in either approach. Therefore, our pipeline has adopted both methods.

**Figure 3.**
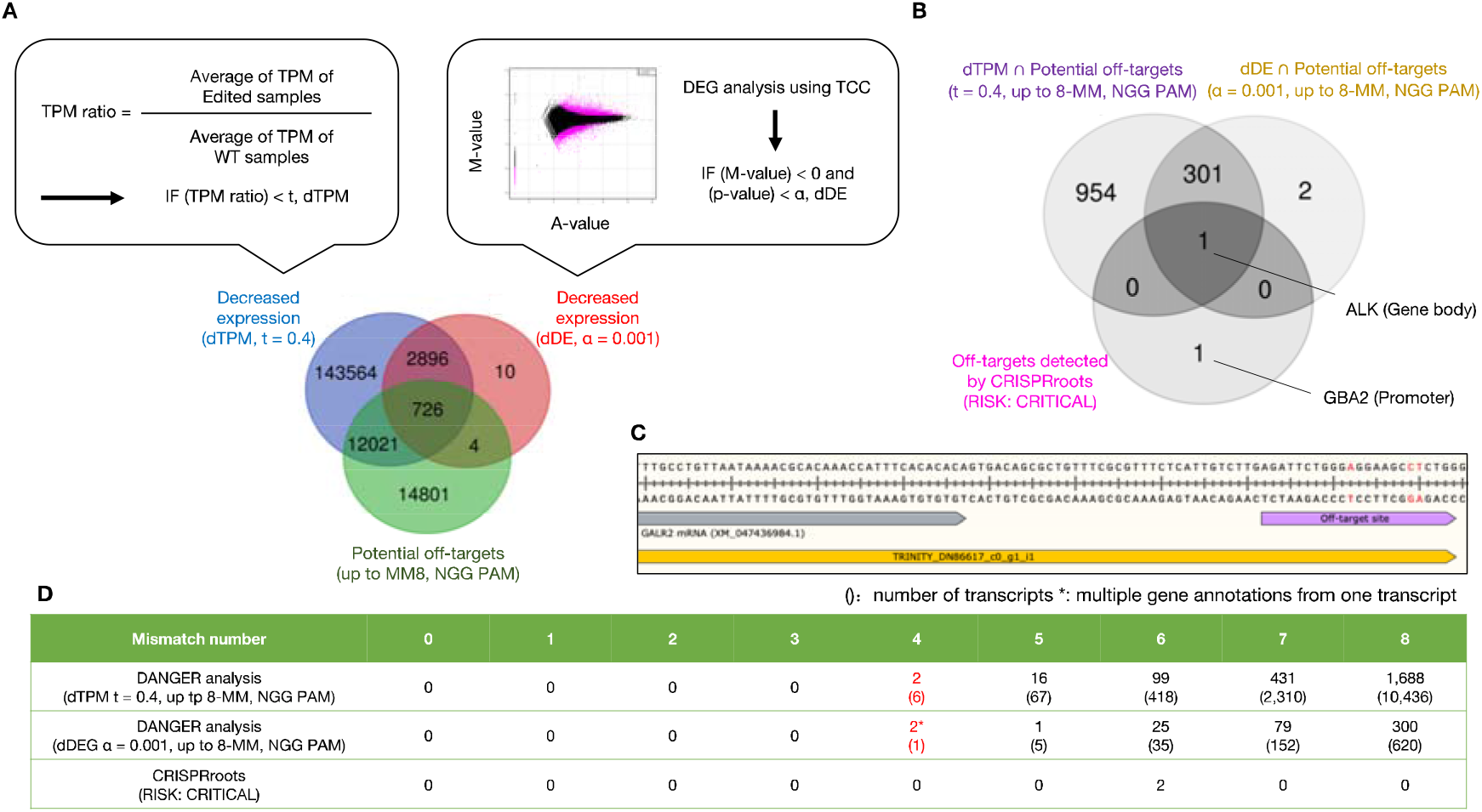
The benchmark for expression analysis methods compared with reference-based RNA-seq analysis using RNA-seq data derived from WT and GRIN2B edited iPSC-derived cortical neurons. A. Comparison of different expression analyses. A Venn diagram comparing the *de novo* transcripts (duplicate counts on a predicted ORF basis), which had potential off-target sites with up to 8 nt mismatches, was detected by the dTPM and dDE approaches. dTPM indicates that the expression is decreased based on the ratio of TPM counts between WT and Edited samples (left callout). dDE means the expression is reduced based on DEG analysis between WT and Edited samples (right callout). B. Comparison of *de novo* transcriptome assembly-based and reference-based analysis on the deleterious off-target detection. A Venn diagram comparing the off-target genes identified from *de novo* transcriptome analysis (dTPM (t = 0.4) and dDE (α = 0.001) approaches) and reference-based RNA-seq analysis (CRISPRroots, "RISK: CRITICAL"). C. Genomic sequence map of off-target located outside of GALR2 mRNA. The sequence is a part of the hg38 reference genome with annotations of GALR2 mRNA (XM_047436984.1) and the *de novo* transcript (TRINITY_DN86617_c0_g1_i1) and an off-target site with three mismatches compared to the on-target sequence. D. Summary of deleterious off-target sites detected by *de novo* transcriptome analysis (dTPM) and reference-based RNA-seq analysis (CRISPRroots, "RISK: CRITICAL"). D. The counts of off-target sites are annotated with genes and classified by mismatch number related to the on-target sequence. The brackets indicate the number of transcripts, including those with and without identified gene annotations. The number of genes and transcripts with ≤4 nt mismatches is colored red.

The Gorodkin and Seemann group previously established the on/off-target assessment pipeline (CRISPRroots)(^44^). They then used STAR(^50^) to perform expression analysis on the same RNA-seq data of GRIN2B using reference-based mapping. They suggested that two off-target sites (ALK: gcAgaCTGGtTGGAAGCaCCNGG, GBA2: cccTTCcGGccGGAAGCGCCNGG) were binding with the Cas9-GRIN2B-REV sgRNA (AGATTCTGGGTGGAAGCGCCNGG)-DNA seed and were linked to downregulated expression (namely, “RISK: CRITICAL”). The definition of off-targets in their study was similar to our idea of deleterious off-targets. Thus, we compared our list of deleterious off-targets in dTPM and dDE with the two off-targets detected using CRISPRroots. As a result, in the DANGER analysis, the off-target of CRISPRroots in the ALK locus was detected by both dTPM (t = 0.4) and dDE (α = 0.001) criteria considering up to eight base mismatches with NGG PAM, whereas the off-target of CRISPRroots in GBA2 was not detected by any criteria (Figure 3B). The absence of GBA2 was understandable because the off-target site on the GBA2 locus was in the promoter region rather than the GBA2 transcript. Additionally, more stringent expression analyses were conducted using dTPM (t = 0.2) and dDE (α = 0.0001). As a result, although 125 off-target genes were detected, neither dTPM nor dDE could identify ALK as an off-target gene (Supplementary Figure S2). This result suggested the possibility of an increase in false negatives when the threshold for detection of expression decreases resulting from off-target effects is overly strict. It is therefore considered appropriate to set t = 0.4 in dTPM and α = 0.001 in dDE. Our *de novo* transcriptome approach focuses solely on the potential off-target genes in the transcribed region of the genome. Additionally, unlike traditional reference-based RNA-seq analysis, this approach can provide novel insights. For example, the off-target search of DANGER analysis detected a transcript with an off-target site downstream GALR2 locus (Figure 3C). The genome database annotation was not the off-target site of the transcribed region (XM_047436984.1). However, the *de novo* transcriptome assembly included the site in the transcript (TRINITY_DN86617_c0_g1_i1). This result indicates that our DANGER analysis pipeline can detect *bona fide* transcripts, which have never been annotated in the reference genome database because of cell-specific transcription, personal genomic variants, and inadequate genomic locus study. Additionally, the *de novo* transcriptome assembly had 260,770 transcript annotations (contigs), which may include transcript variants that partially came from allelic heterogeneity. The annotation size of DANGER analysis is about ten times larger than that of CRISPRroots, whose gene annotations were about 25k. DANGER Analysis was expected to make a larger off-target dataset of transcribed regions using the transcript-aware annotations compared to reference-based analysis. Although the detection range of our DANGER analysis is limited to the transcribed region, our pipeline using dTPM and dDE detected 13,237 and 813 off-target sites with zero to eight mismatches in identified and unidentified transcripts, respectively. 2,236 and 407 gene-annotated off-target sites with four to eight mismatches, respectively. In contrast, genome-wide and reference-based RNA-seq analysis (CRISPRroots) features only two off-target sites in genes with six mismatches. There was a large discrepancy in the detection number of off-targets between DANGER Analysis and CRISPRroots. We demonstrated that the DANGER analysis could generate a comprehensive and factual record of off-target sites (Figure 3D).

Finally, our pipeline evaluated the phenotypic risk of deleterious off-targets. Various studies have used Gene Ontology (GO) analysis to assess phenotypic effects(^20,23,31–34^). However, this study also considered off-target frequency because deleterious off-targets with many mismatches were expected to occur(^5,46^). Thus, our pipeline counted off-target genes associated with specific GO terms per mismatch number (Figure 4A) and then combined the results with the mismatch effect by calculating the D-index per GO term (Figure 4B; see Materials and methods). The D-index is calculated by multiplying the number of genes containing a GO term for each mismatch number by a decreasing exponential function with the mismatch number as the exponent. This approach allows us to consider the number of genes hit by the GO terms and the number of mismatches. Moreover, it can suppress the influence of the number of genes hit when the mismatch number is large. The formula can prioritize evaluating the number of genes hit when the mismatch number is small. With these two characteristics, the D-index represents a unique off-target metric that considers the impact on phenotype. The sum of the D-index value (total D-index) in detected GO terms was 6,228 (N= 9,896) (Figure 4C, Supplementary Table S2) in the GRIN2B dataset. The DANGER analysis was used to evaluate the phenotypic risk at the GO level using RNA-seq of the human GRIN2B dataset data without any reference genomes.

**Figure 4.**
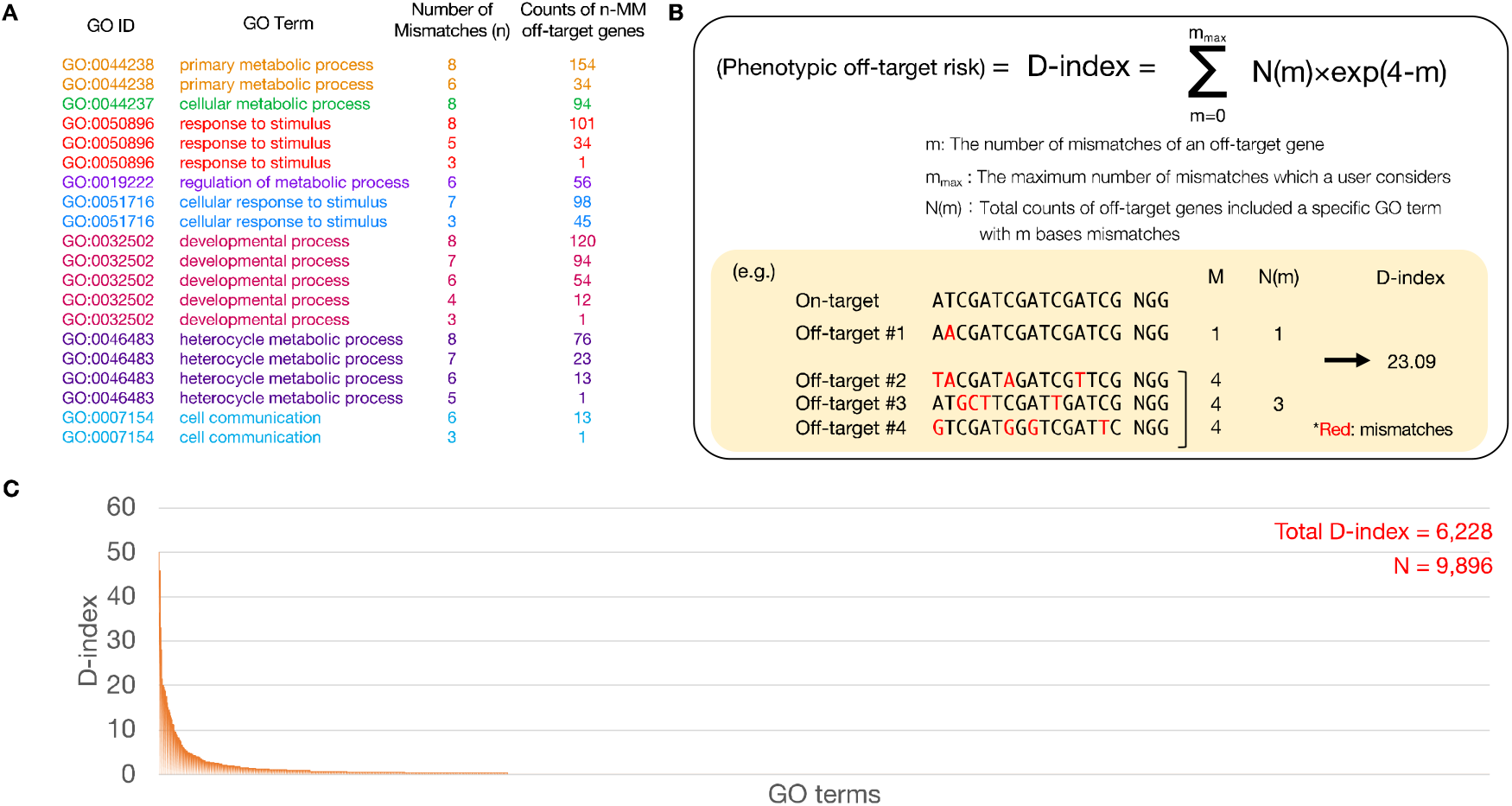
The result of risk assessment in DANGER analysis using RNA-seq data derived from WT and GRIN2B edited iPSC-derived cortical neurons. A. An example of the annotation table for DANGER analysis. The table includes GO ID, GO term, number of mismatches (n), and the counts of n-MM off-target genes belonging to a specific GO term. B. The formula for phenotypic off-target risk (D-index). An example of the calculation is shown on a yellow background. C. Distribution of the D-index of each GO term (orange). The sum of all D-indexes and the number of D-indices (N) were labeled on the top right.

### Evaluation of D-index and optimization of DANGER analysis

Using the D-index, we quantified the phenotypic impact from off-targets at the GO term level. However, a different threshold must be set for each GO term when evaluating the statistical significance of the D-index values because GO is a loosely hierarchical annotation concept, and ‘parent’ terms appear as annotations of various genes even if there is less relationship with off-targets. To address this issue, we implemented a permutation test system to estimate the significance threshold for each GO term (Figure 5). In this system, after randomly shuffling the expression profile and off-target profile, we repeat the process of calculating a meaningless D-index (named pseudo-D-index) by applying the D-index formula 100 times. We then create a null distribution from the 100 pseudo-D-index values for each GO term and define a Significant D-index as an originally obtained D-index value that exceeds the (1 – L) ×100 % confidence interval of the null distribution. The methodology and the threshold allow us to extract only the D-indices of the GO terms suspected of having an off-target-associated impact on the phenotype.

**Figure 5.**
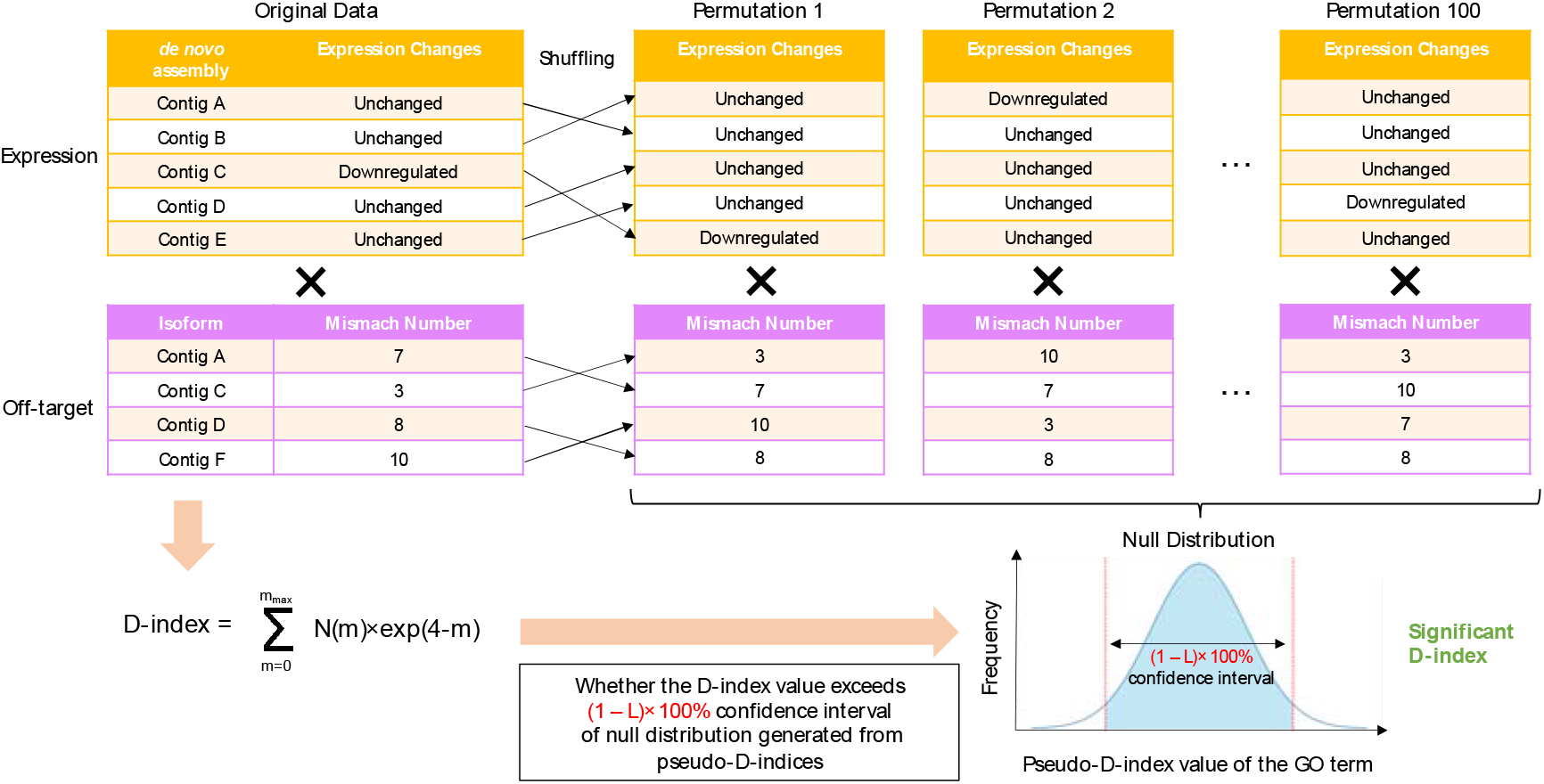
A scheme for permutation testing to evaluate the validity of the D-index. The thin black arrow indicates the manipulation of rearranging values from the original expression and off-target profile to the permutation data. The black cross represents the computation for applying the D-index formula to the above expression profile and the below off-target profile data. The workflow is shown as the orange allows.

Moreover, we devised a scheme to assess the validity of the permutation test system (Supplementary Figure S3). In the validation scheme, followed by generating several different pseudo-D indices, the number of the newly generated pseudo-D indices exceeding the threshold of the previous null distribution is counted. We defined the ratio of the count to the total number of D-indices as the false detection rate in the permutation test. We used the evaluation scheme to verify how the false detection rate changes under various conditions in DANGER Analysis. No significant influence was observed from the threshold of the expression analysis (t, α) or the conditions of off-target search (Supplementary Figure S4A-C). We confirmed L=1E-15 confidence interval threshold reduced the false detection rate by more than half in the case of L=5E-1. Here, we named the condition of dTPM with high false positives (up to 11-MM NRR PAM, t=0.4, L=5E-1) as ’Approximate dTPM,’ the condition of dTPM with low false positives (up to 8-MM NGG PAM, t=0.4, L=1E-15) as ’Optimized dTPM,’ and the condition of dDE with low false positives (up to 8-MM NGG PAM, α = 0.001, L=1E-15) as ’Optimized dDE.’ When comparing the three conditions, the optimized dDE showed the lowest false detection rate (Figure 6A). Next, we calculated the total number of D-indices, significant D-indices, and true significant D-indices (Expected True D-indices) estimated from the False Detection Rate for these three conditions. The total number of D-indices was more than twice as high for dTPM as for dDE, while the number of Expected True D-indices was less than half for dTPM compared to dDE (Figure 6B). The result indicates that the dTPM criterion is a ’sharply-narrowing-down’ approach used for initial screening, while dDE is a ’meticulously trimming’ approach used for a rigorous selection process. Generally, the dDE approach is recommended for use in human RNA-seq data because the criterion is expected to present fewer false detections in significant D-indices. However, in model organisms and non-model organisms with less comprehensive GO annotations than humans, the dDE approach may not yield a sufficient D-index list for the evaluation. Thus, the dTPM approach, which can obtain more D-indices, is expected to be more effective in RNA-seq data from less characterized organisms than humans. We evaluated the consistency of D-indices and Significant D-indices detected by both optimized dTPM and optimized dDE. The concordance of each D-index and Significant D-index was approximately 52% and 1.3%, respectively (Figure 6C), which can be attributed to the fact that optimized dTPM considers more than five times the gene-annotated off-target sites compared to optimized dDE (Figure 3D). However, the significant D-indices commonly detected by optimized dTPM and optimized dDE corresponded to the top 16 significant D-indices in dTPM (Figure 3C-D). The result suggested that the value of the D-index not only served as an indicator of the phenotypic impact from off-targets but could also be an indicator of the strength of its consistency. Thus, it is recommended to conduct follow-up analyses focusing mainly on the top-ranking D-indices in optimized dTPM.

**Figure 6.**
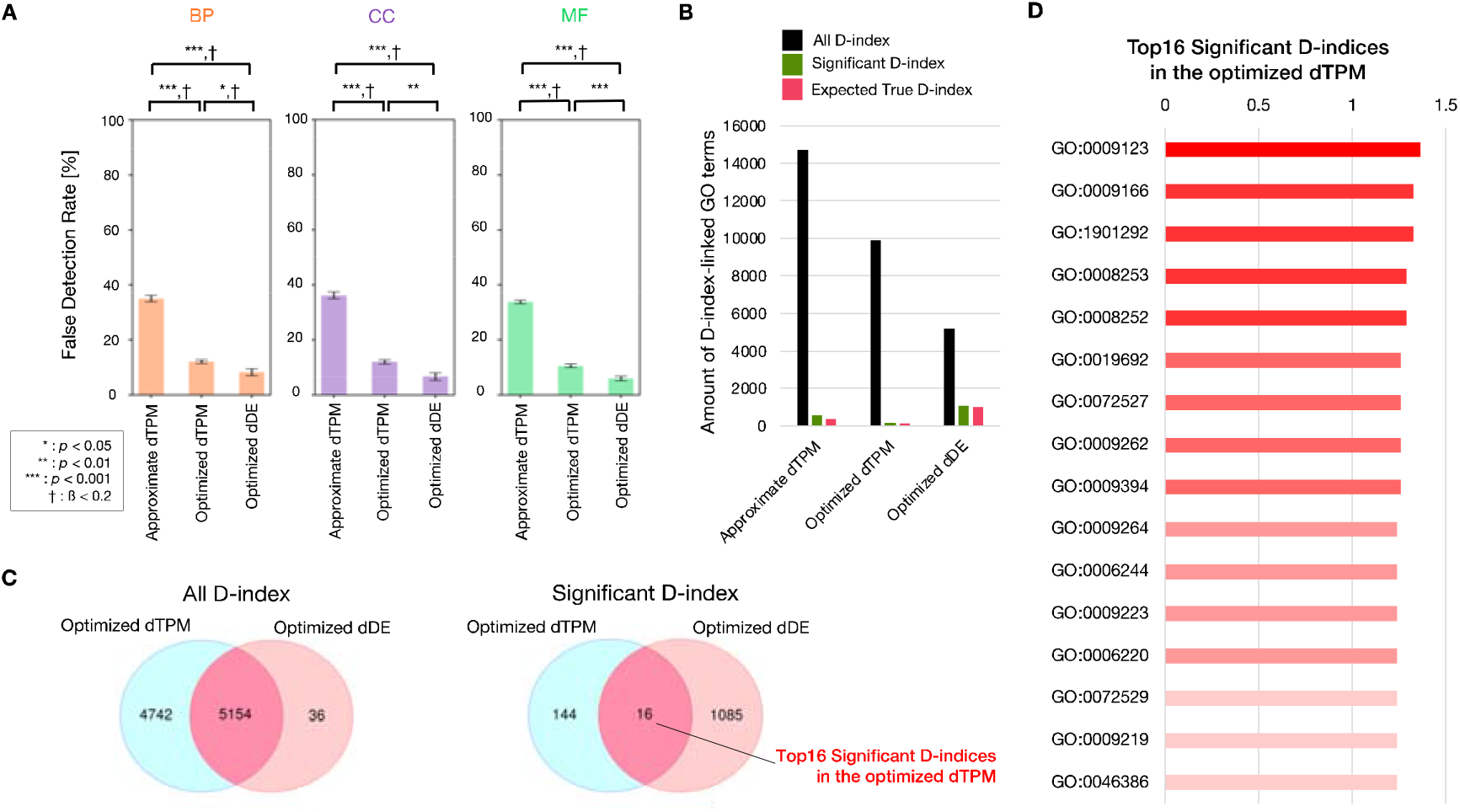
Evaluation of permutation test for DANGER analysis and comparison between dTPM and dDE. A. Comparison of false detection rates among approximate dTPM (up to 11- MM NRR PAM, t=0.4, L=5E-1), optimized dTPM (up to 8-MM NGG PAM, t=0.4, L=1E-15), and optimized dDE (up to 8-MM NGG PAM, α = 0.001, L=1E-15) in GO categories. BP, CC, and MF indicate GO categories of Biological Process, Cellular Component, and Molecular Function, respectively. Error bars represent SEM; asterisk indicates the statistical significance of two-sided Welch’s t-test; cross indicates statistical power (1-ß) >0.8. Mean ± s.d. of n = 10 permutation data set. B. Comparison of amount of GO terms of all D-index, significant D-index, and expected true D-index among approximate dTPM (up to 11-MM NRR PAM, t=0.4, L=5E-1), optimized dTPM (up to 8-MM NGG PAM, t=0.4, L=1E-15), and optimized dDE (up to 8-MM NGG PAM, α = 0.001, L=1E-15), respectively. C. Comparison of D-index and significant D-index between optimized dTPM and optimized dDE. A Venn diagram comparing the counts of D-index and significant D-index between optimized dTPM and optimized dDE. D. The list of the top 16 significant D-indices in the optimized dTPM. The D-index values are indicated by bar graphs adjacent to the GO terms.

### Assessment of CRISPR-Cas9 on/off-target using DANGER analysis for RNA-seq data from *in vivo* tissue of zebrafish

We performed DANGER analysis using the RNA-seq data from human cells edited by one of the Cas9n-sgRNAs for benchmarking with the previous method (CRISPRroots) and optimized parameters for the DANGER analysis in the previous sections. Next, we investigated whether DANGER analysis could be used to analyze the RNA-seq data from the non-human tissue that had been *in vivo* edited with a single Cas9 nuclease-sgRNA, a more common experimental design for genome editing. Thus, we downloaded and analyzed RNA-seq data from the park7 dataset derived from the zebrafish brain, with and without indels at the park7 locus(^20^). Our DANGER analysis successfully built a *de novo* transcriptome assembly with 90.9% complete BUSCO genes and detected on-target sequences in the two transcripts (Figure 7A). Moreover, DANGER analysis revealed a significant downregulation of the transcript with Cas9-sgRNA on-target sequence in the expression quantification (Figure 7B). The original report on the park7 dataset reported downregulated park7 mRNA in the park7 mutant using RNA-seq analysis(^20^). This result implied that *de novo* transcriptome assembly and the following expression quantification of our DANGER analysis could generate reliable data consistent with the outcome of standard RNA-seq analysis using a reference genome. Consequently, 19,314 potential off-target sites in all transcripts were detected by optimized dTPM, which were then defined as deleterious off-targets considering the expression profile. There were 4,668 and 70 deleterious off-target sites on all and gene-annotated transcripts, respectively (Figure 7C). The park7 result had no deleterious off-target effects with ≤ 4 mismatches, which meant that frequent off-targeting was not expected in mRNA-transcribed regions. The detected rate of gene-annotated transcripts to all transcripts was more than ten times less than that of optimized dTPM using human GRIN2B (Figure 3D and Figure 7C). The fewer annotations resulted from the poor gene database of zebrafish in comparison with that of humans. Next, our pipeline estimated the D-index per GO term to quantify phenotypic risk. The total D-index was 51 (N=636) (Figure 7D and Supplementary Table S5), which was less than that of human GRIN2B due to fewer annotations and off-target genes. Finally, we validated the D-indices using the permutation test as the same procedure in the last section. Only five significant D-indices were detected (Figure 7E) because the analysis considered only 70 deleterious off-target sites in gene-annotated transcripts. As discussed in the previous section, the park7 analysis has empirically demonstrated that the initially considered number of genes and the total number of D-indices can be small in organisms other than humans. The screening approach of optimized dTPM allows for the acquisition of significant D-indices in poorly annotated data sets.

**Figure 7.**
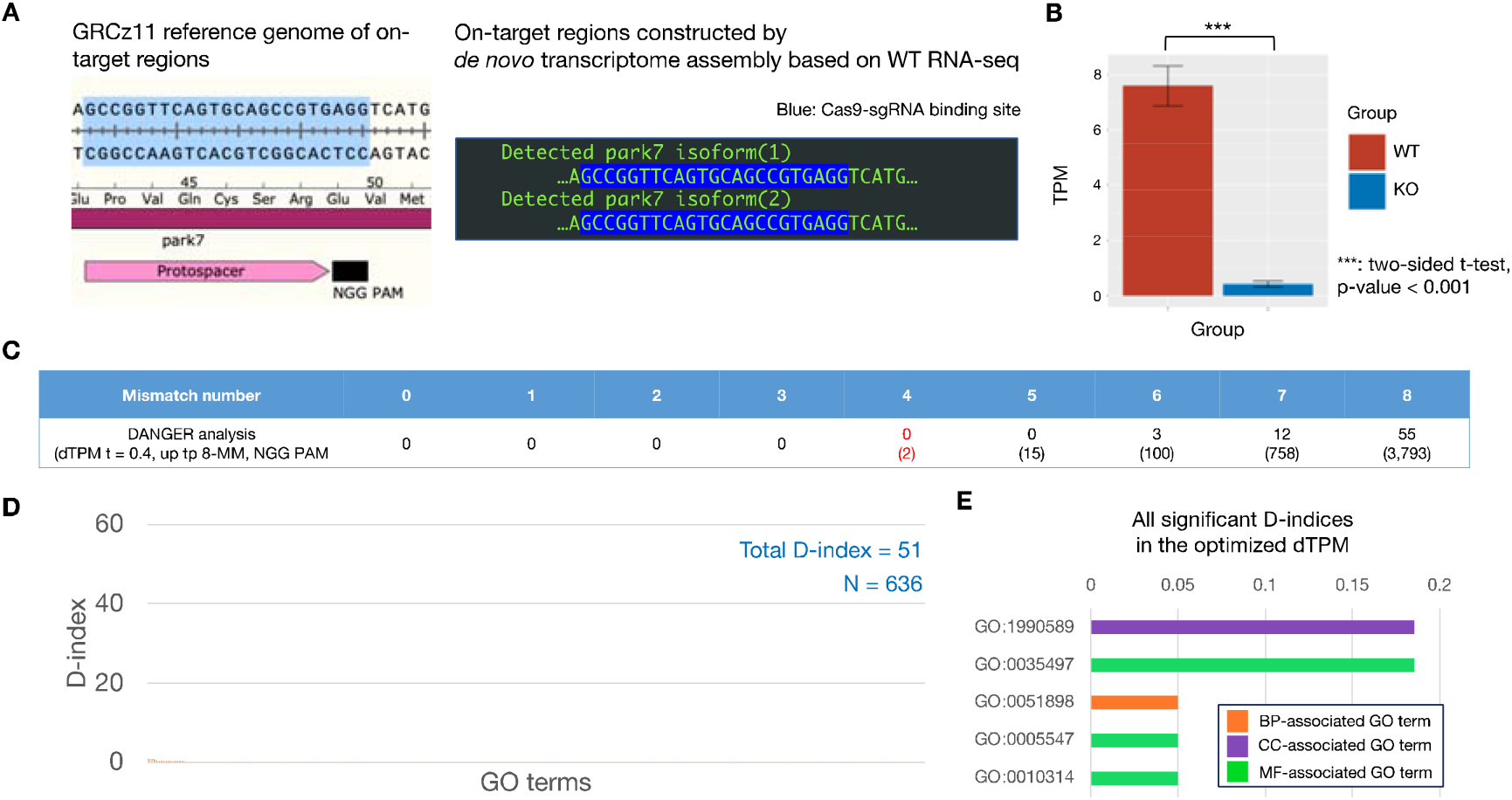
DANGER analysis result using RNA-seq data derived from WT and park7 (dj1) Edited brains of *Danio rerio*. A. Comparison between the GRCz11 reference genome and transcript sequence constructed by *de novo* assembly of RNA-seq samples derived from WT brain on park7 on-target region. The on-target region of the GRCz11 reference genome is illustrated with annotations of the park7 CDS, the protospacer, and the NGG PAM sequence of SpCas9. The detected park7 isoforms (1-2) are lined up in green in the black box. The Cas9-sgRNA binding sites are highlighted in blue. B. Comparison of TPM values of park7. The TPM was measured from WT and Edited RNA-seq samples (Each n=3); data were expressed as the means±SEM. *** p-value < 0.001 of two-sided Welch’s t-test. C. The gene counts are classified by mismatch number related to the on-target sequence. The brackets indicate the number of transcripts, including those with and without identified gene annotations. The number of genes and transcripts with ≤4 nt mismatches is colored red. D. Distribution of the D-index of each GO term associated with Biological Process (orange). The sum of all D-indices and the number of D-indices (N) is labeled on the top right. E. The list of all significant D-indices in the optimized dTPM. The D-index values are indicated by bar graphs adjacent to the GO terms. The colors of the bar indicate GO categories belonging to the GO terms.

### Comparison of phenotypic risks in the GRIN2B and Park7 datasets

In this study, we evaluated the phenotypic risks associated with off-target transcripts using the D-index. The number of significant D-indices of the GRIN2B result was approximately 32- fold larger than that of the park7 result in the optimized dTPM. Furthermore, the DANGER analysis found off-target genes with four mismatches in the GRIN2B dataset, which is common in genome-wide off-target studies such as GUIDE-seq and Digenome-seq(^24,26^) and numerous deleterious off-target sites on transcript sequences annotated with genes. Thus, Cas9n-GRIN2B-REV sgRNA may have side effects on the phenotype of differentiated human iPSC. A follow-up study is required to assess the edited GRIN2B LOF clones using WGS or alternative genome-wide methods such as GUIDE-seq(^24^). Researchers can identify some clarified points in the future using the result table of the significant D-index (Supplementary Table S3-4). For example, the GO term “nucleoside monophosphate metabolic process” (GO ID: GO:0009123) recorded a top significant D-index for the GRIN2B result of optimized dTPM (Supplementary Table S3). The GO terms of "cell differentiation“ (GO ID: GO:0030154), “cell population proliferation” (GO ID: GO:0008283), and “cell cycle” (GO ID: GO:0007049) were considered as the phenotype of GRIN2B knock-out in the previous study(^23^). However, these GO terms were listed in the significant D-indices of optimized dDE (Supplementary Table S4, Table 2). The previous study showed that the gene expression changes of these GO terms resulted from on-target editing of the GRIN2B locus(^23^). However, the results of the DANGER analysis suggested that off-target editing of additional genes belonging to GO:0030154, GO:0008283, and GO:0007049 partially contributed to expression changes. Follow-up studies should include off-target gene analysis of the associated off-targets with the GO terms. On the other hand, The GO terms of "central nervous system development “ (GO ID: GO:0007417), “brain development” (GO ID: GO:0007420), “cell division” (GO ID: GO:0051301), and “chromosome segregation” (GO ID: GO:0007059) were not listed in the significant D-indices of optimized dTPM and optimized dDE, which suggested the GO terms were obvious phenotypes in the GRIN2B knock-out cells. Therefore, DANGER analysis would help reach a reasonable conclusion in genome editing studies.

**Table 2.**
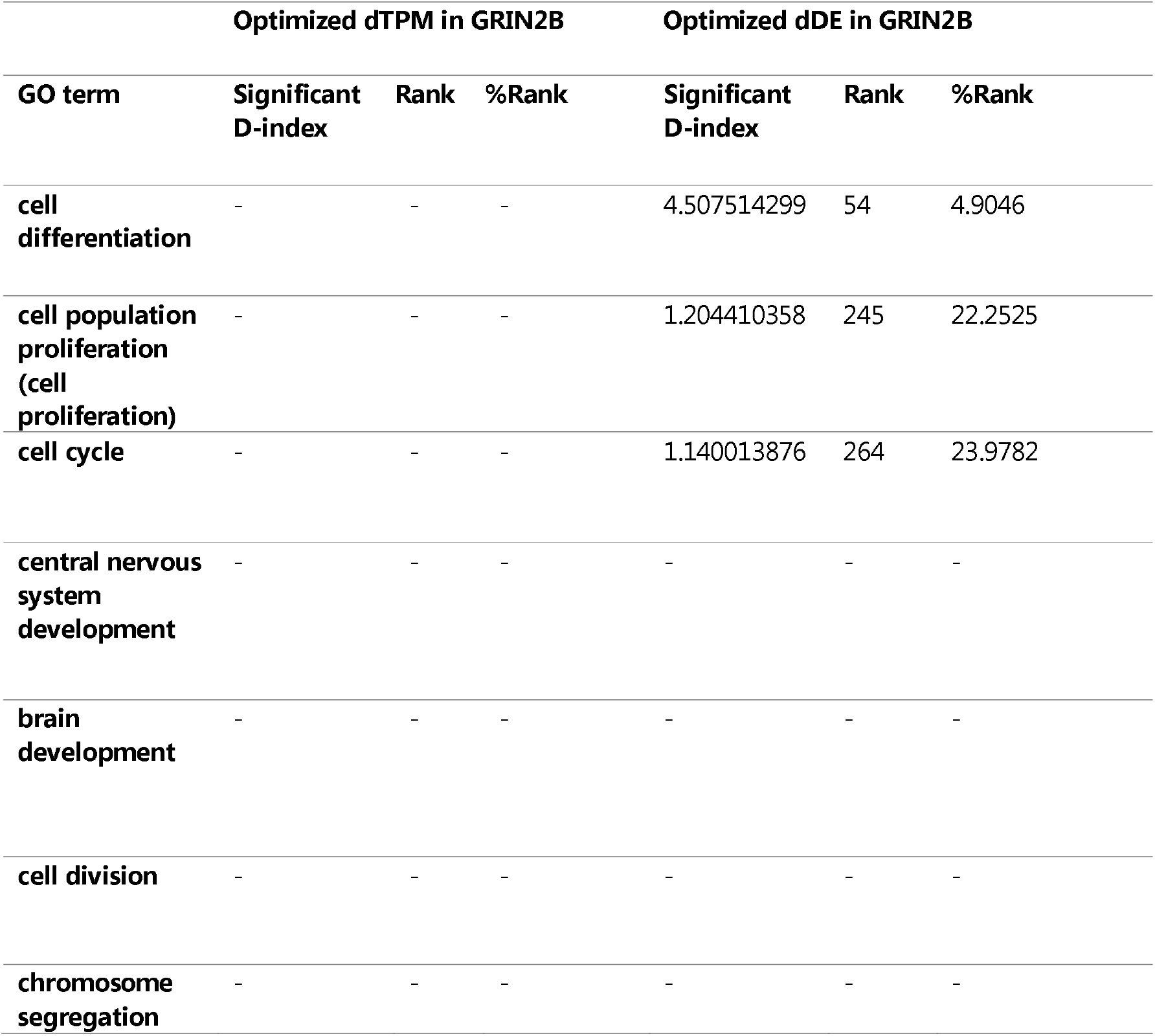
D-index summary of GO terms related to the enrichment analysis of previous GRIN2B research.

### Limitations

In this section, we discuss the major limitations of the proposed pipeline. First, our method depends on the quality of *de novo* transcriptome assembly using Trinity. Pair-end RNA-seq data with sufficient length and read number must be used to guarantee high-quality assembly (see Material and Methods). If we fail to build an exemplary assembly, producing reliable data for the following analyses, such as on/off-target analysis and expression profiles, becomes difficult. Second, the annotation analysis step in our pipeline may fail to annotate transcripts adequately due to limited information from databases on genes, transcripts, proteins, and gene function. When researchers apply our pipeline to RNA-seq data from organisms with limited genomic knowledge and evidence, we recommend using a database of a model organism with a strong genomic relationship with the organism being analyzed. DANGER analysis can analyze genome-edited samples without a reference genome, but studies of a related model organism with a well-annotated genome are still required. As a third limitation, DANGER analysis cannot strictly distinguish the effects of modifications to on-target genes on other genes from the impacts of off-target gene modifications. Of course, an on-target gene is excluded from the DANGER analysis. Still, it is difficult to distinguish the influence by considering off-target genes whose expression is controlled by the on-target genes. Such an evaluation can only be elucidated through protein interaction and genome analysis conducted using more specialized knowledge by researchers in each field. Therefore, it is appropriate to complete the follow-up studies mentioned in the previous section using a comprehensive analysis. Traditional studies have discussed results based on RNA-seq analysis under the premise that they are solely derived from the effects of on-target gene modifications. However, our DANGER analysis contradicts this assumption, sounding an alarm about the necessity for more specialized investigations and providing the off-target gene information needed for such follow-up analyses. Additionally, gene network analyses using found off-target genes can help users exclude false detection if the target organism of DANGER Analysis is a model organism whose gene database is well established.

### Possibility of DANGER Analysis as a Simplified Screening Tool, Contributing to More Rigorous Reference-based Phenotypic Risk Assessment

In this study, we developed DANGER analysis as an initial screening tool for maximizing the evaluation of risks to phenotypes. Meanwhile, it is conceivable that we will also need a more rigorous evaluation system for assessing risks to phenotypes. In constructing such an evaluation system, it is believed that a system utilizing information other than RNA-seq data, such as reanalysis of sample genomes by resequencing the genomic DNA rather than *de novo* transcriptome assembly, would be appropriate. Although our DANGER analysis is a tool that only takes RNA-seq data as input to ensure convenience, there is room to apply the partial algorithm (association between off-target genes and GO terms, phenotype risk calculations using the D-index) into such a rigorous reference-based evaluation system for phenotype risks. We believe there is a high possibility that DANGER analysis could become a foundational presence in this new field of phenotype risk assessment.

### DANGER analysis Provides a New Perception of the Conventional Genome Editing Process in Medicine, Agriculture, and Biological Research

We demonstrated our DANGER analysis pipeline, as it allows for (i) the detection of potential DNA on/off-target sites in the mRNA-transcribed region on the genome using RNA-seq data, (ii) evaluation of phenotypic effects by deleterious off-target sites based on the evidence provided by gene expression changes, and (iii) quantification of the phenotypic risk at the GO term level, without a reference genome. Thus, DANGER analysis can be performed on various organisms, personal human genomes, and atypical genomes created by diseases and viruses(^28^). The CRISPRroots is expected to be only effective in samples with high similarity to the well-characterized reference genome. In general, DANGER analysis holds superiority over CRISPRroots in terms of versatility. We believe that the perception resulting from our DANGER analysis has not been observed in the conventional scheme for genome editing. We illustrate a new scheme using DANGER analysis in organisms (I) with and (II) without a reference genome (Figure 8). For example, (I) model organisms such as humans have a reference genome whose information has been generally used for potential on/off-target searches and post-analysis. However, the reference genome is a representative genome and not the personal genome of the patient or cell lines. Reference-based genome editing does not consider unique single nucleotide polymorphisms (SNPs) or spontaneous genomic rearrangements. DANGER analysis can supply a personal transcriptome-based on/off-target profile to ensure the phenotypic risk of unexpected off-target mutations. The new workflow of genome editing would be helpful for *ex vivo* gene therapy and cancer research because the genome of a cancer cell is generally characterized by widespread somatic genomic rearrangements(^28^). (II) An organism whose genome has never been comprehensively sequenced and well characterized is not considered a reasonable subject for genome editing, as site-specific genome editing is guaranteed without a reference genome. Some groups, however, have used incomplete genomic information to construct mutants of non-model organisms(^51,52^) that have never been well-characterized in genomics. Such a conventional scheme is hit-or-miss due to the risk of erroneous knock-out phenotypes in combination with off-targeting of other genes. DANGER analysis can provide transcriptome phenotype-aware on-/off-target profiles as well as sequence information of the expressed genes. This information can be used for the safer design of genome editing, which enables the optimized design of CRISPR editing by repeatedly looping back through genome-editing experiments without a reference genome. Furthermore, the DANGER analysis devised in this study employs a simple algorithm based on mismatch count for identifying off-target candidates. The type of algorithm has also been utilized in off-target investigations for other programmable nucleases such as CRISPR-Cas12a, TALEN, and ZFN. DANGER analysis is open-source and freely adjustable. Thus, the algorithm of this pipeline could be repurposed for the analysis of various genome editing systems beyond the CRISPR-Cas9 system. Moreover, it is also possible to enhance the specificity of DANGER Analysis for CRISPR-Cas9 by incorporating a CRISPR-Cas9 specific off-target scoring algorithms. We believe that the DANGER analysis pipeline will expand the scope of genomic studies and industrial applications using genome editing.

**Figure 8.**
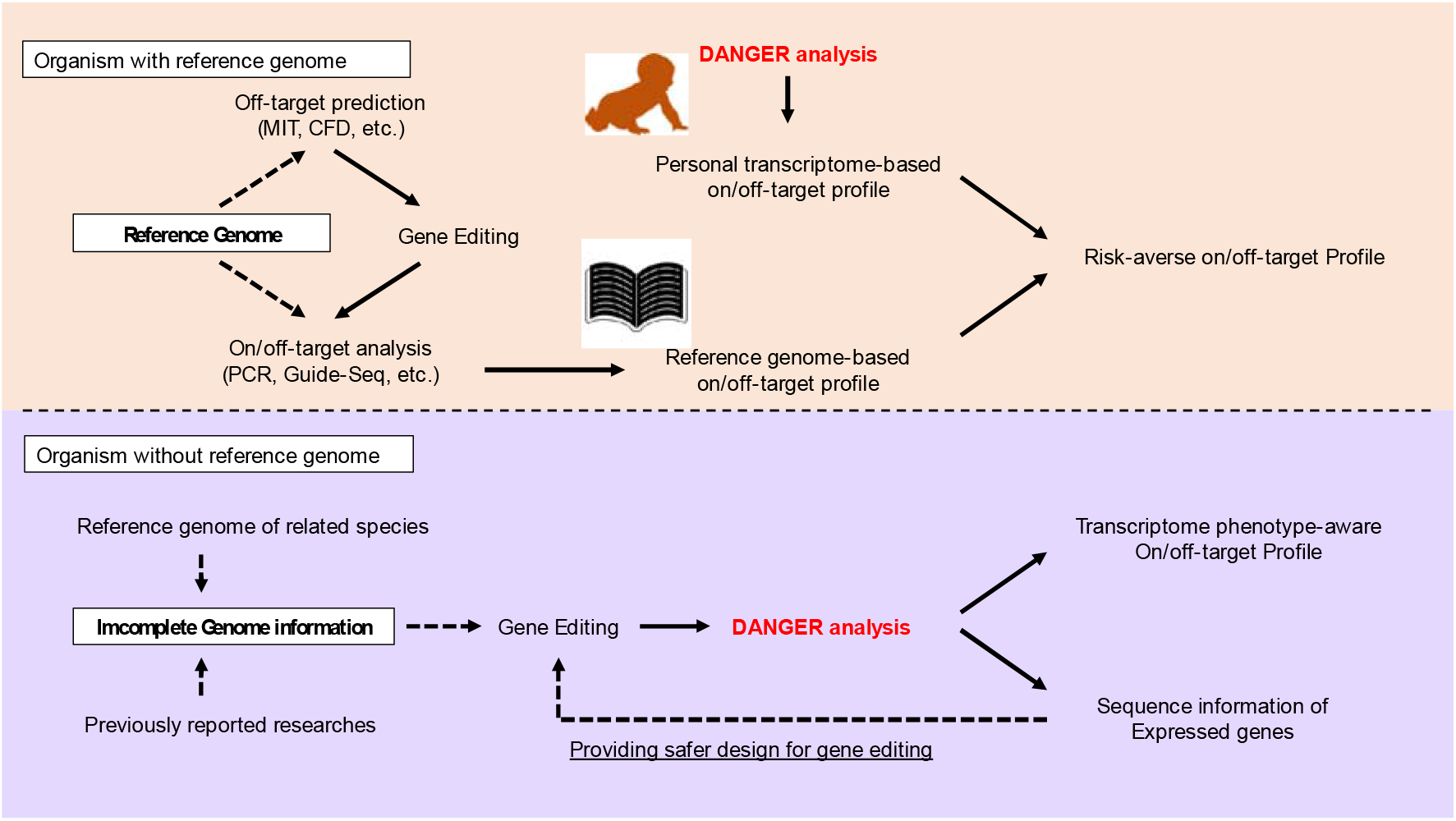
Our proposal for the usage of DANGER-analysis in organisms with and without a reference genome. The workflow is shown as black arrows. The dotted black arrows indicate the front of the arrow and refer to the arrow base information. The image of the book is from TogoTV (© 2016 DBCLS TogoTV, CC-BY-4.0, https://creativecommons.org/licenses/by/4.0/).

## DATA AVAILABILITY

The datasets were derived from the following public domain resources: https://www.ncbi.nlm.nih.gov/geo. The analyses were performed using DANGER analysis version 1.0. The Script for the DANGER analysis pipeline is available at https://github.com/KazukiNakamae/DANGER_analysis. In addition, the software provides a tutorial on reproducing the results presented in this article on the Readme page. The Docker image of DANGER_analysis is also available at https://hub.docker.com/repository/docker/kazukinakamae/dangeranalysis/general.

## SUPPLEMENTARY DATA

Supplemental_Files - zip file

Supplementary Data are available at *Bioinformatics Advances* online.

## AUTHOR CONTRIBUTIONS

Kazuki Nakamae: Conceptualization, Software, Formal analysis, Methodology, Validation, Visualization, Writing of the original draft. Hidemasa Bono: Supervision, Writing – Review, and Editing.

## ACKNOWLEDGEMENTS

We would like to thank all the laboratory members at Hiroshima University, PtBio Inc., and RIKEN for their valuable comments. We deeply appreciate Dr. Ryo Nozu for providing us with a supplemental script (https://github.com/RyoNozu/Sequence_editor). We would like to thank Editage (www.editage.com) for English language editing.

## REFERENCES

1. Wiedenheft, B., Sternberg, S.H., and Doudna, J.A. (2012). RNA-guided genetic silencing systems in bacteria and archaea. Nature 482, 331–338. 10.1038/nature10886.

2. Terns, M.P., and Terns, R.M. (2011). CRISPR-Based Adaptive Immune Systems. Curr Opin Microbiol 14, 321–327. 10.1016/j.mib.2011.03.005.

3. Jinek, M., East, A., Cheng, A., Lin, S., Ma, E., and Doudna, J. (2013). RNA-programmed genome editing in human cells. eLife 2, e00471. 10.7554/eLife.00471.

4. Gillmore, J.D., Gane, E., Taubel, J., Kao, J., Fontana, M., Maitland, M.L., Seitzer, J., O’Connell, D., Walsh, K.R., Wood, K., et al. (2021). CRISPR-Cas9 In Vivo Gene Editing for Transthyretin Amyloidosis. New England Journal of Medicine 385, 493–502. 10.1056/NEJMoa2107454.

5. Hsu, P.D., Scott, D.A., Weinstein, J.A., Ran, F.A., Konermann, S., Agarwala, V., Li, Y., Fine, E.J., Wu, X., Shalem, O., et al. (2013). DNA targeting specificity of RNA-guided Cas9 nucleases. Nat Biotechnol 31, 827–832. 10.1038/nbt.2647.

6. Bassett, A.R., Tibbit, C., Ponting, C.P., and Liu, J.-L. (2013). Highly Efficient Targeted Mutagenesis of Drosophila with the CRISPR/Cas9 System. Cell Rep 4, 220–228. 10.1016/j.celrep.2013.06.020.

7. Shirai, Y., Piulachs, M.-D., Belles, X., and Daimon, T. (2022). DIPA-CRISPR is a simple and accessible method for insect gene editing. Cell Reports Methods 2, 100215. 10.1016/j.crmeth.2022.100215.

8. Nymark, M., Sharma, A.K., Sparstad, T., Bones, A.M., and Winge, P. (2016). A CRISPR/Cas9 system adapted for gene editing in marine algae. Sci Rep 6, 24951. 10.1038/srep24951.

9. Yoshimitsu, Y., Abe, J., and Harayama, S. (2018). Cas9-guide RNA ribonucleoprotein-induced genome editing in the industrial green alga Coccomyxa sp. strain KJ. Biotechnology for Biofuels 11, 326. 10.1186/s13068-018-1327-1.

10. Jiang, W., Bikard, D., Cox, D., Zhang, F., and Marraffini, L.A. (2013). CRISPR-assisted editing of bacterial genomes. Nat Biotechnol 31, 233–239. 10.1038/nbt.2508.

11. Jinek, M., Chylinski, K., Fonfara, I., Hauer, M., Doudna, J.A., and Charpentier, E. (2012). A Programmable Dual-RNA–Guided DNA Endonuclease in Adaptive Bacterial Immunity. Science 337, 816–821. 10.1126/science.1225829.

12. Cong, L., Ran, F.A., Cox, D., Lin, S., Barretto, R., Habib, N., Hsu, P.D., Wu, X., Jiang, W., Marraffini, L.A., et al. (2013). Multiplex Genome Engineering Using CRISPR/Cas Systems. Science 339, 819–823. 10.1126/science.1231143.

13. Mojica, F.J.M., Díez-Villaseñor, C., García-Martínez, J., and Almendros, C. (2009). Short motif sequences determine the targets of the prokaryotic CRISPR defence system. Microbiology 155, 733–740. 10.1099/mic.0.023960-0.

14. Fu, Y., Sander, J.D., Reyon, D., Cascio, V.M., and Joung, J.K. (2014). Improving CRISPR-Cas nuclease specificity using truncated guide RNAs. Nat Biotechnol 32, 279–284. 10.1038/nbt.2808.

15. Qi, L.S., Larson, M.H., Gilbert, L.A., Doudna, J.A., Weissman, J.S., Arkin, A.P., and Lim, W.A. (2013). Repurposing CRISPR as an RNA-Guided Platform for Sequence-Specific Control of Gene Expression. Cell 152, 1173–1183. 10.1016/j.cell.2013.02.022.

16. Komor, A.C., Kim, Y.B., Packer, M.S., Zuris, J.A., and Liu, D.R. (2016). Programmable editing of a target base in genomic DNA without double-stranded DNA cleavage. Nature 533, 420–424. 10.1038/nature17946.

17. Nishida, K., Arazoe, T., Yachie, N., Banno, S., Kakimoto, M., Tabata, M., Mochizuki, M., Miyabe, A., Araki, M., Hara, K.Y., et al. (2016). Targeted nucleotide editing using hybrid prokaryotic and vertebrate adaptive immune systems. Science 353, aaf8729. 10.1126/science.aaf8729.

18. Anzalone, A.V., Randolph, P.B., Davis, J.R., Sousa, A.A., Koblan, L.W., Levy, J.M., Chen, P.J., Wilson, C., Newby, G.A., Raguram, A., et al. (2019). Search-and-replace genome editing without double-strand breaks or donor DNA. Nature 576, 149–157. 10.1038/s41586-019-1711-4.

19. Gagnon, J.A., Valen, E., Thyme, S.B., Huang, P., Ahkmetova, L., Pauli, A., Montague, T.G., Zimmerman, S., Richter, C., and Schier, A.F. (2014). Efficient Mutagenesis by Cas9 Protein-Mediated Oligonucleotide Insertion and Large-Scale Assessment of Single-Guide RNAs. PLOS ONE 9, e98186. 10.1371/journal.pone.0098186.

20. Hughes, G.L., Lones, M.A., Bedder, M., Currie, P.D., Smith, S.L., and Pownall, M.E. (2020). Machine learning discriminates a movement disorder in a zebrafish model of Parkinson’s disease. Dis Model Mech 13, dmm045815. 10.1242/dmm.045815.

21. Liu, D., Awazu, A., Sakuma, T., Yamamoto, T., and Sakamoto, N. (2019). Establishment of knockout adult sea urchins by using a CRISPR-Cas9 system. Development, Growth & Differentiation 61, 378–388. 10.1111/dgd.12624.

22. Koike-Yusa, H., Li, Y., Tan, E.-P., Velasco-Herrera, M.D.C., and Yusa, K. (2014). Genome-wide recessive genetic screening in mammalian cells with a lentiviral CRISPR-guide RNA library. Nat Biotechnol 32, 267–273. 10.1038/nbt.2800.

23. Bell, S., Maussion, G., Jefri, M., Peng, H., Theroux, J.-F., Silveira, H., Soubannier, V., Wu, H., Hu, P., Galat, E., et al. (2018). Disruption of GRIN2B Impairs Differentiation in Human Neurons. Stem Cell Reports 11, 183–196. 10.1016/j.stemcr.2018.05.018.

24. Tsai, S.Q., Zheng, Z., Nguyen, N.T., Liebers, M., Topkar, V.V., Thapar, V., Wyvekens, N., Khayter, C., Iafrate, A.J., Le, L.P., et al. (2015). GUIDE-seq enables genome-wide profiling of off-target cleavage by CRISPR-Cas nucleases. Nat Biotechnol 33, 187–197. 10.1038/nbt.3117.

25. Luo, X., He, Y., Zhang, C., He, X., Yan, L., Li, M., Hu, T., Hu, Y., Jiang, J., Meng, X., et al. (2019). Trio deep-sequencing does not reveal unexpected off-target and on-target mutations in Cas9-edited rhesus monkeys. Nat Commun 10, 5525. 10.1038/s41467-019- 13481-y.

26. Kim, D., Bae, S., Park, J., Kim, E., Kim, S., Yu, H.R., Hwang, J., Kim, J.-I., and Kim, J.-S. (2015). Digenome-seq: genome-wide profiling of CRISPR-Cas9 off-target effects in human cells. Nat Methods 12, 237–243. 10.1038/nmeth.3284.

27. Levy, S., Sutton, G., Ng, P.C., Feuk, L., Halpern, A.L., Walenz, B.P., Axelrod, N., Huang, J., Kirkness, E.F., Denisov, G., et al. (2007). The Diploid Genome Sequence of an Individual Human. PLOS Biology 5, e254. 10.1371/journal.pbio.0050254.

28. Vogelstein, B., Papadopoulos, N., Velculescu, V.E., Zhou, S., Diaz, L.A., and Kinzler, K.W. (2013). Cancer Genome Landscapes. Science 339, 1546–1558. 10.1126/science.1235122.

29. Ashburner, M., Ball, C.A., Blake, J.A., Botstein, D., Butler, H., Cherry, J.M., Davis, A.P., Dolinski, K., Dwight, S.S., Eppig, J.T., et al. (2000). Gene Ontology: tool for the unification of biology. Nat Genet 25, 25–29. 10.1038/75556.

30. The Gene Ontology resource: enriching a GOld mine (2020). Nucleic Acids Res 49, D325–D334. 10.1093/nar/gkaa1113.

31. Tamura, K., and Bono, H. (2022). Meta-Analysis of RNA Sequencing Data of Arabidopsis and Rice under Hypoxia. Life 12, 1079. 10.3390/life12071079.

32. Toga, K., Yokoi, K., and Bono, H. (2022). Meta-Analysis of Transcriptomes in Insects Showing Density-Dependent Polyphenism. Insects 13, 864. 10.3390/insects13100864.

33. Bono, H. (2021). Meta-Analysis of Oxidative Transcriptomes in Insects. Antioxidants 10, 345. 10.3390/antiox10030345.

34. Suzuki, T., Ono, Y., and Bono, H. (2021). Comparison of Oxidative and Hypoxic Stress Responsive Genes from Meta-Analysis of Public Transcriptomes. Biomedicines 9, 1830. 10.3390/biomedicines9121830.

35. Subramanian, A., Tamayo, P., Mootha, V.K., Mukherjee, S., Ebert, B.L., Gillette, M.A., Paulovich, A., Pomeroy, S.L., Golub, T.R., Lander, E.S., et al. (2005). Gene set enrichment analysis: A knowledge-based approach for interpreting genome-wide expression profiles. Proceedings of the National Academy of Sciences 102, 15545–15550. 10.1073/pnas.0506580102.

36. Hölzer, M., and Marz, M. (2019). De novo transcriptome assembly: A comprehensive cross-species comparison of short-read RNA-Seq assemblers. Gigascience 8, giz039. 10.1093/gigascience/giz039.

37. Lipka, A., Paukszto, L., Majewska, M., Jastrzebski, J.P., Panasiewicz, G., and Szafranska, B. (2019). De novo characterization of placental transcriptome in the Eurasian beaver (Castor fiber L.). Funct Integr Genomics 19, 421–435. 10.1007/s10142-019-00663-6.

38. Grabherr, M.G., Haas, B.J., Yassour, M., Levin, J.Z., Thompson, D.A., Amit, I., Adiconis, X., Fan, L., Raychowdhury, R., Zeng, Q., et al. (2011). Full-length transcriptome assembly from RNA-Seq data without a reference genome. Nat Biotechnol 29, 644–652. 10.1038/nbt.1883.

39. Khudyakov, J.I., Champagne, C.D., Meneghetti, L.M., and Crocker, D.E. (2017). Blubber transcriptome response to acute stress axis activation involves transient changes in adipogenesis and lipolysis in a fasting-adapted marine mammal. Sci Rep 7, 42110. 10.1038/srep42110.

40. Nespolo, R.F., Gaitan-Espitia, J.D., Quintero-Galvis, J.F., Fernandez, F.V., Silva, A.X., Molina, C., Storey, K.B., and Bozinovic, F. (2018). A functional transcriptomic analysis in the relict marsupial Dromiciops gliroides reveals adaptive regulation of protective functions during hibernation. Molecular Ecology 27, 4489–4500. 10.1111/mec.14876.

41. Martin, M. (2011). Cutadapt removes adapter sequences from high-throughput sequencing reads. EMBnet.journal 17, 10–12. 10.14806/ej.17.1.200.

42. Manni, M., Berkeley, M.R., Seppey, M., Simão, F.A., and Zdobnov, E.M. (2021). BUSCO Update: Novel and Streamlined Workflows along with Broader and Deeper Phylogenetic Coverage for Scoring of Eukaryotic, Prokaryotic, and Viral Genomes. Mol Biol Evol 38, 4647–4654. 10.1093/molbev/msab199.

43. Jacquin, A.L.S., Odom, D.T., and Lukk, M. (2019). Crisflash: open-source software to generate CRISPR guide RNAs against genomes annotated with individual variation. Bioinformatics 35, 3146–3147. 10.1093/bioinformatics/btz019.

44. Corsi, G.I., Gadekar, V.P., Gorodkin, J., and Seemann, S.E. (2021). CRISPRroots: on-and off-target assessment of RNA-seq data in CRISPR–Cas9 edited cells. Nucleic Acids Res 50, e20. 10.1093/nar/gkab1131.

45. Sun, J., Nishiyama, T., Shimizu, K., and Kadota, K. (2013). TCC: an R package for comparing tag count data with robust normalization strategies. BMC Bioinformatics 14, 219. 10.1186/1471-2105-14-219.

46. Fu, R., He, W., Dou, J., Villarreal, O.D., Bedford, E., Wang, H., Hou, C., Zhang, L., Wang, Y., Ma, D., et al. (2022). Systematic decomposition of sequence determinants governing CRISPR/Cas9 specificity. Nat Commun 13, 474. 10.1038/s41467-022-28028- x.

47. Kurosaki, T., Popp, M.W., and Maquat, L.E. (2019). Quality and quantity control of gene expression by nonsense-mediated mRNA decay. Nat Rev Mol Cell Biol 20, 406–420. 10.1038/s41580-019-0126-2.

48. Srivastava, A., Malik, L., Sarkar, H., Zakeri, M., Almodaresi, F., Soneson, C., Love, M.I., Kingsford, C., and Patro, R. (2020). Alignment and mapping methodology influence transcript abundance estimation. Genome Biol 21, 239. 10.1186/s13059-020-02151-8.

49. Wagner, G.P., Kin, K., and Lynch, V.J. (2012). Measurement of mRNA abundance using RNA-seq data: RPKM measure is inconsistent among samples. Theory Biosci. 131, 281–285. 10.1007/s12064-012-0162-3.

50. Dobin, A., Davis, C.A., Schlesinger, F., Drenkow, J., Zaleski, C., Jha, S., Batut, P., Chaisson, M., and Gingeras, T.R. (2013). STAR: ultrafast universal RNA-seq aligner. Bioinformatics 29, 15–21. 10.1093/bioinformatics/bts635.

51. Chang, K.S., Kim, J., Park, H., Hong, S.-J., Lee, C.-G., and Jin, E. (2020). Enhanced lipid productivity in AGP knockout marine microalga Tetraselmis sp. using a DNA-free CRISPR-Cas9 RNP method. Bioresource Technology 303, 122932. 10.1016/j.biortech.2020.122932.

52. Sun, Q., Lin, L., Liu, D., Wu, D., Fang, Y., Wu, J., and Wang, Y. (2018). CRISPR/Cas9-Mediated Multiplex Genome Editing of the BnWRKY11 and BnWRKY70 Genes in Brassica napus L. International Journal of Molecular Sciences 19, 2716. 10.3390/ijms19092716.

