## Supplemental Information for "DANGER ANALYSIS: RISK-AVERSE ON/OFF-TARGET ASSESSMENT FOR CRISPR EDITING WITHOUT A REFERENCE GENOME"

#### Supplementary Information

##### Contents

1. Supplementary Figures . . . . . . . . . . . . . . . . . . . . . . . . . . . . . . . . . . . . . . . . . . . . . . . . . . .. . . . 2
2. Supplementary Tables . . . . . . . . . . . . . . . . . . . . . . . . . . . . . . . . . . . . . . . . . . . . . . . . . . . . . . 7

### 1 Supplementary Figures

A

| Mismatch number | 0 | 1 | 2 | 3 | 4 | 5 | 6 | 7 | 8 |
|---|---|---|---|---|---|---|---|---|---|
| dTPM (t = 0.4) only | 0 | 0 | 0 | 0 | 5 | 62 | 383 | 2161 | 9817 |
| dDE (α = 0.001) only | 0 | 0 | 0 | 0 | 0 | 0 | 0 | 3 | 1 |
| dTPM (t = 0.4) ∩ dDE (α = 0.001) | 0 | 0 | 0 | 0 | 1 | 5 | 35 | 149 | 619 |

B

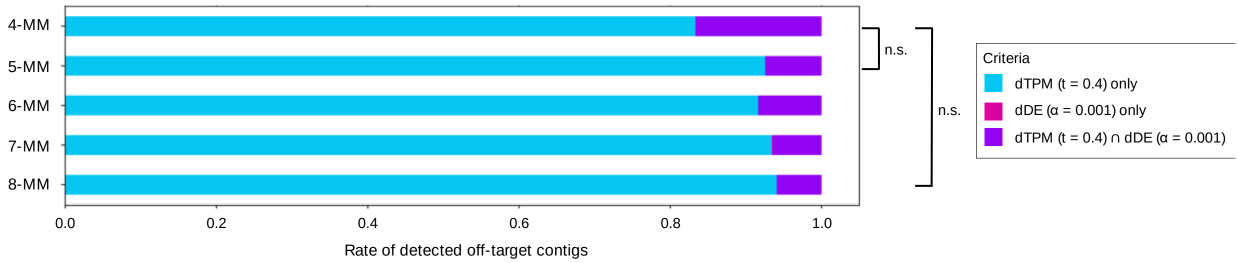


Fig. S1: A comparison of off-target mismatches detected by dTPM and dDE, categorized by the number of mismatches. A. Comparison of mismatch counts for off-targets detected solely by dTPM (dTPM only), solely by dDE (dDE only), or commonly detected by both dTPM (t=0.4) and dDE ($\alpha$ = 0.001) (dTPM ∩ dDE). B. stacked bar plot of mismatch counts for off-targets detected by "dTPM only," "dDE only," or "dTPM ∩ dDE." The black line represents Fisher's exact test for the ratio of off-targets detected by "dTPM only" to those detected by "dTPM ∩ dDE." 'n.s.' stands for 'not significant.'

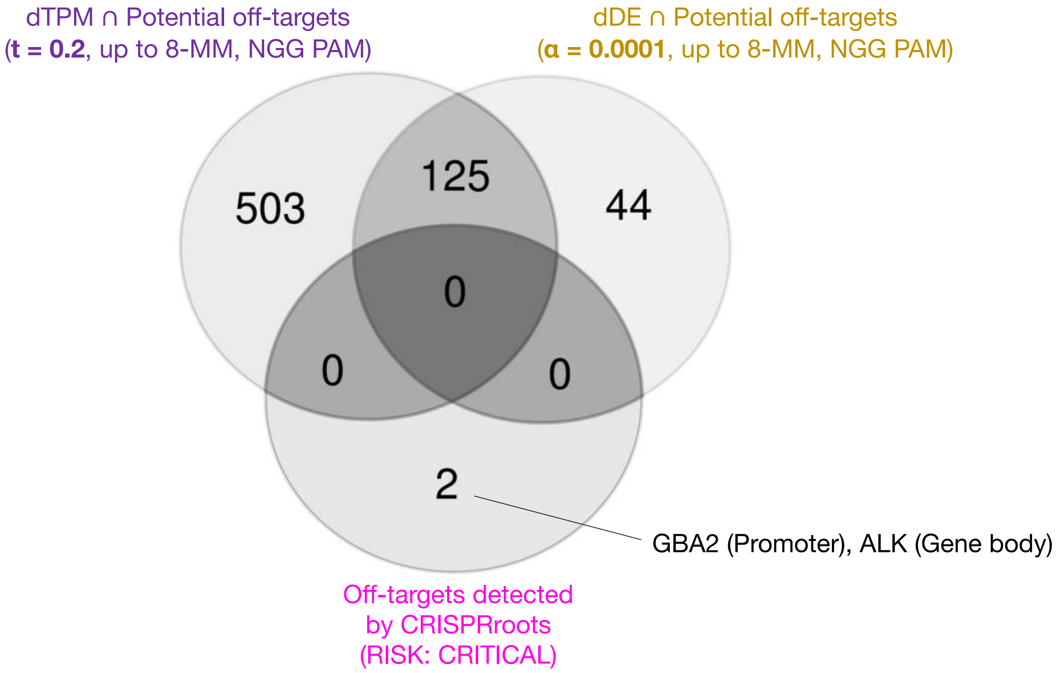


Fig. S2: Comparison of *de novo* transcriptome assembly-based and reference-based analysis on the deleterious off-target detection. A Venn diagram comparing the off-target genes identified from *de novo* transcriptome analysis (dTPM ($t = 0.2$) and dDE($\alpha = 0.0001$) approaches) and reference-based RNA-seq analysis (CRISPRroots, "RISK: CRITICAL").

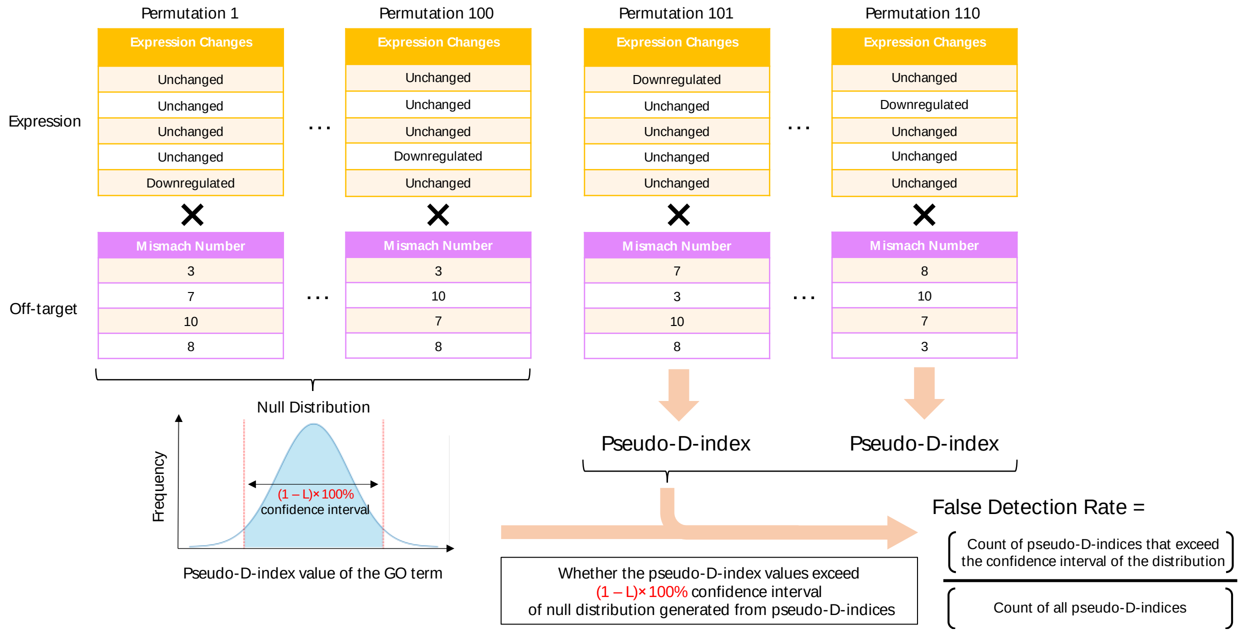


Fig. S3: A scheme for assessment of permutation testing for estimating false detection rate. The black cross represents the computation for applying the D-index formula to the data of the above expression profile and the below off-target profile for generating a pseudo-D-index. The workflow is shown as the orange allows.

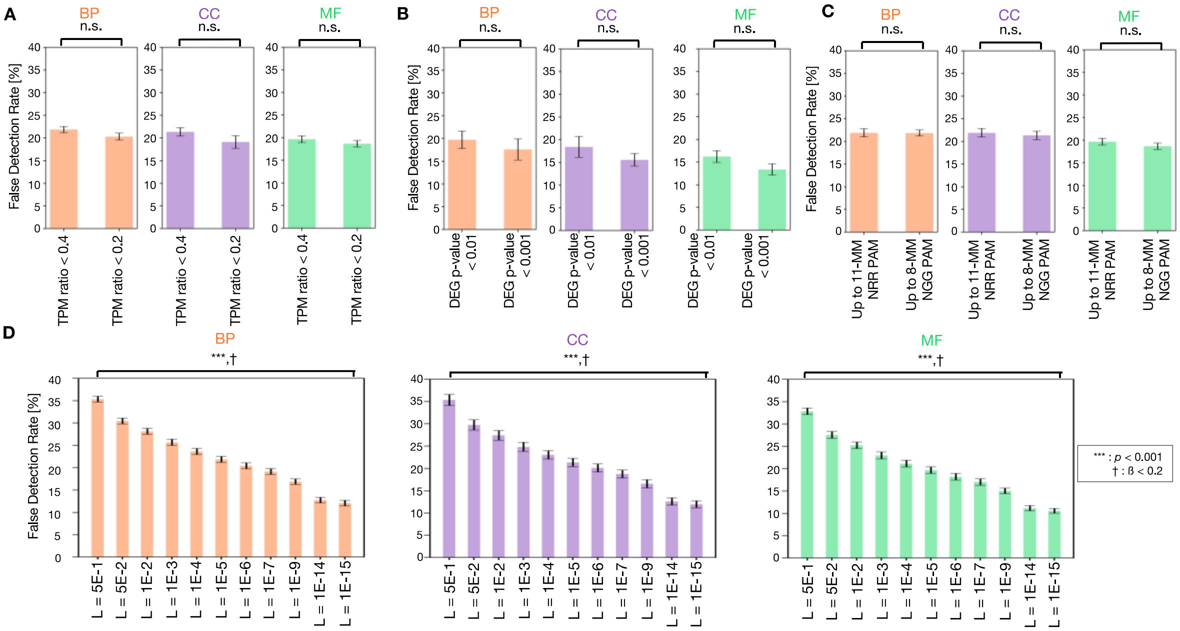


Fig. S4: False detection rate with varying parameters for DANGER analysis. A. False detection rate with varying the threshold (t) for dTPM in GO categories. BP, CC, and MF indicate GO categories of Biological Process, Cellular Component, and Molecular Function, respectively. All off-target analyses considered mismatches ranging from 1 to 8 bases, and the PAM was set as NGG. Also, the confidence interval threshold for the permutation test was set at (1 – L) ×100% = (1 – 1E-05) ×100%. Error bars represent SEM. 'n.s.' stands for 'not significant.' Mean ± s.d. of n = 10 permutation data set. B. False detection rate with varying the threshold (α) for dDE in GO categories. BP, CC, and MF indicate GO categories of Biological Process, Cellular Component, and Molecular Function, respectively. All off-target analyses considered mismatches ranging from 1 to 8 bases, and the PAM was set as NGG. Also, the confidence interval threshold for the permutation test was set at (1 – L) ×100% = (1 – 1E-05) ×100%. Error bars represent SEM. 'n.s.' stands for 'not significant.' Mean ± s.d. of n = 10 permutation data set. C. False detection rate with varying the condition for off-target analyses in GO categories. BP, CC, and MF indicate GO categories of Biological Process, Cellular Component, and Molecular Function, respectively. All expression analyses were conducted using dTPM (t= 0.4). Also, the confidence interval threshold for the permutation test was set at (1 – L) ×100% = (1 – 1E-05) ×100%. Error bars represent SEM.

'n.s.' stands for 'not significant.' Mean ± s.d. of n = 10 permutation data set. D. False detection rate with varying the confidence interval threshold for the permutation test in GO categories. BP, CC, and MF indicate GO categories of Biological Process, Cellular Component, and Molecular Function, respectively. All expression analyses were conducted using dTPM (t= 0.4). Also, all off-target analyses considered mismatches ranging from 1 to 8 bases, and the PAM was set as NGG. Error bars represent SEM. The asterisk indicates the statistical significance of the two-sided Welch's t-test; the cross indicates statistical power (1-ß) >0.8. Mean ± s.d. of n = 10 permutation data set.

#### 2 Supplementary Tables

Tab. S1: A summary of off-target mismatches detected by GUIDE-seq (Tsai *et al.*, 2015), categorized by the number of mismatches.

Tab. S2: D-index table of GO terms based on the result of DANGER analysis with optimized dTPM using RNA-seq data derived from WT and GRIN2B edited iPSC-derived cortical neurons.

Tab. S3: The significant D-index table of GO terms based on the result of DANGER analysis with optimized dTPM using RNA-seq data derived from WT and GRIN2B edited iPSC-derived cortical neurons.

Tab. S4: The significant D-index table of GO terms based on the result of DANGER analysis with optimized dDE using RNA-seq data derived from WT and GRIN2B edited iPSC-derived cortical neurons.

Tab. S5: The D-index table of GO terms based on the result of DANGER analysis with optimized dTPM using RNA-seq data derived from WT and park7 (which encodes DJ-1) edited brains of *Danio rerio*.
