## Supplementary figures and images for "DANGER ANALYSIS: RISK-AVERSE ON/OFF-TARGET ASSESSMENT FOR CRISPR EDITING WITHOUT A REFERENCE GENOME"

### Supplemental Figure 1

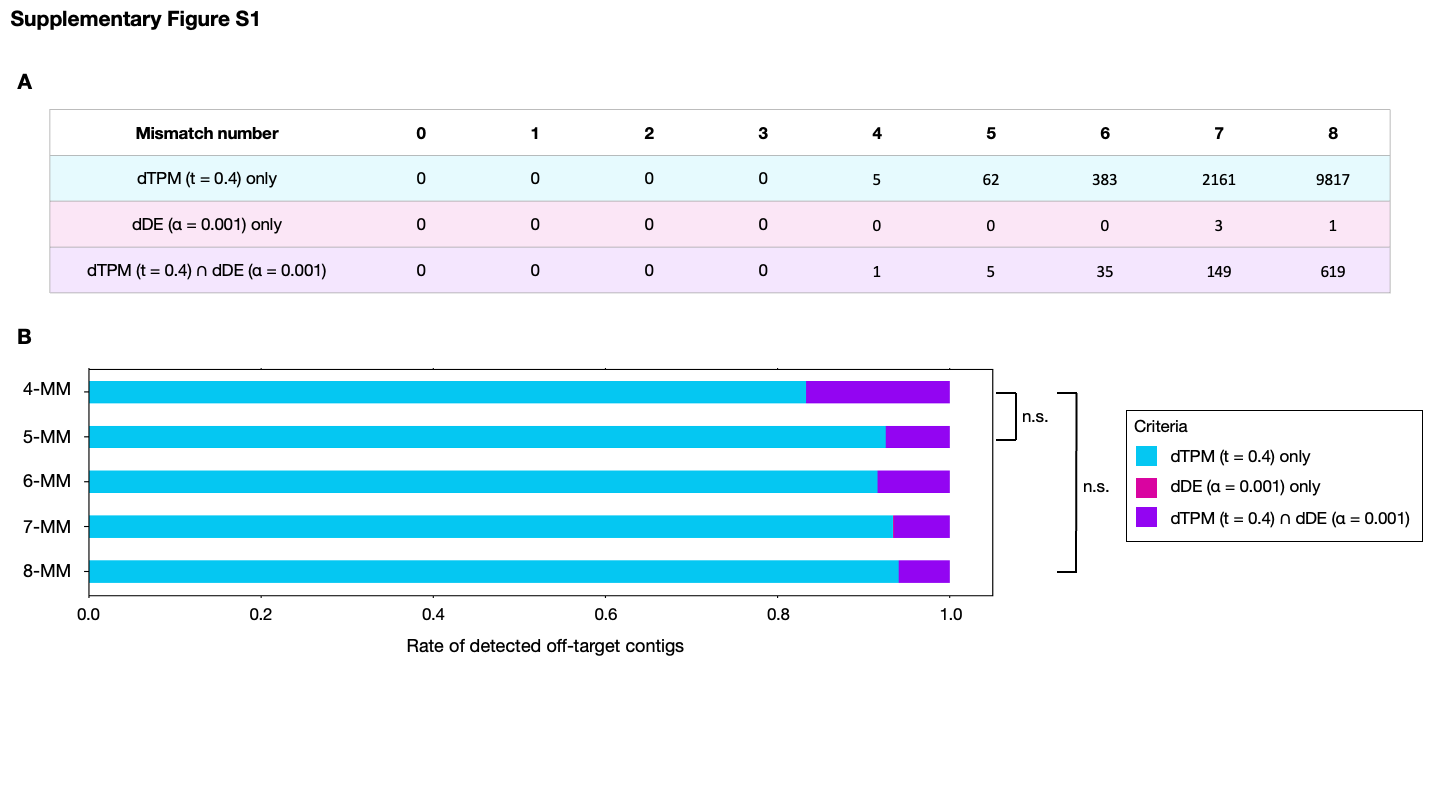

### Supplemental Figure 2

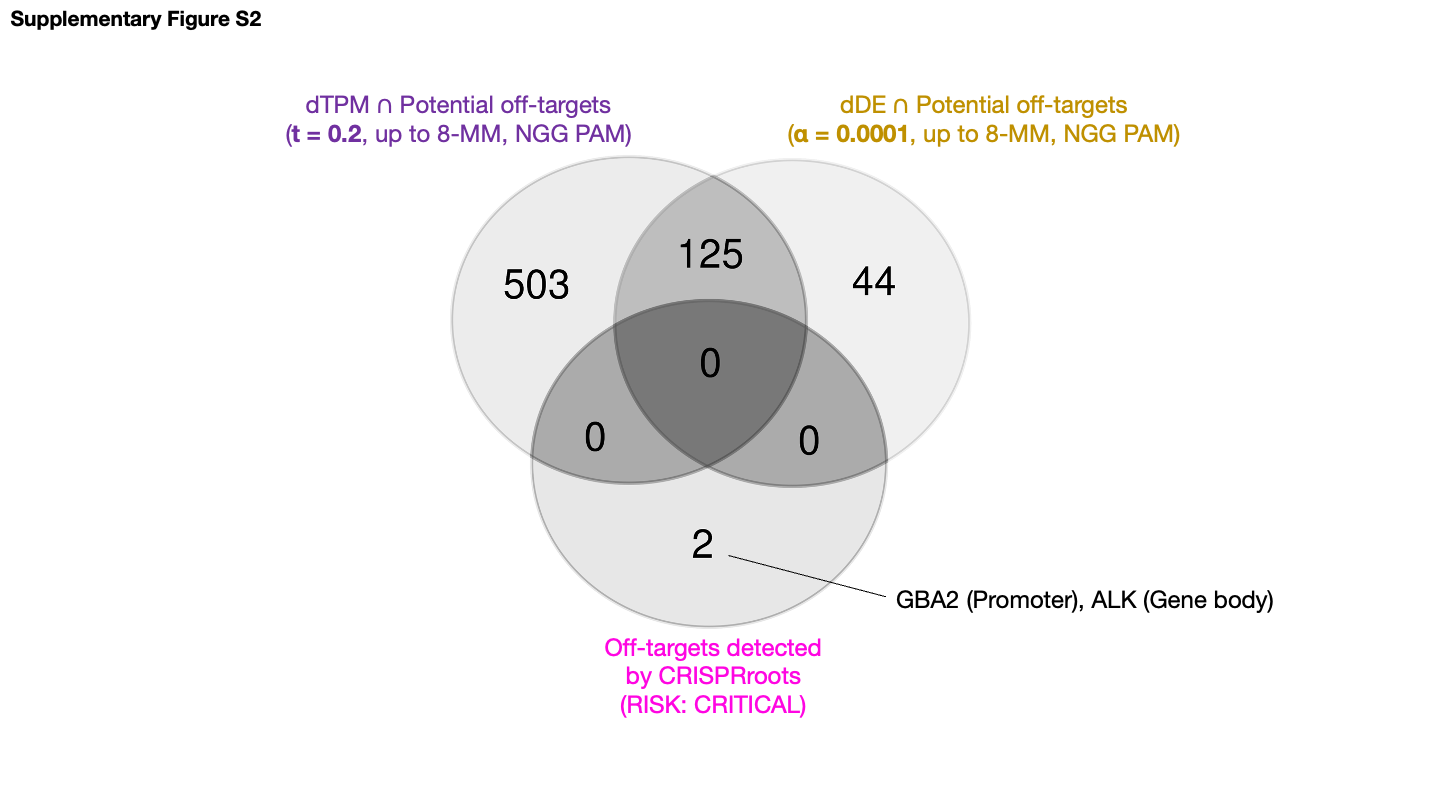

### Supplemental Figure 3

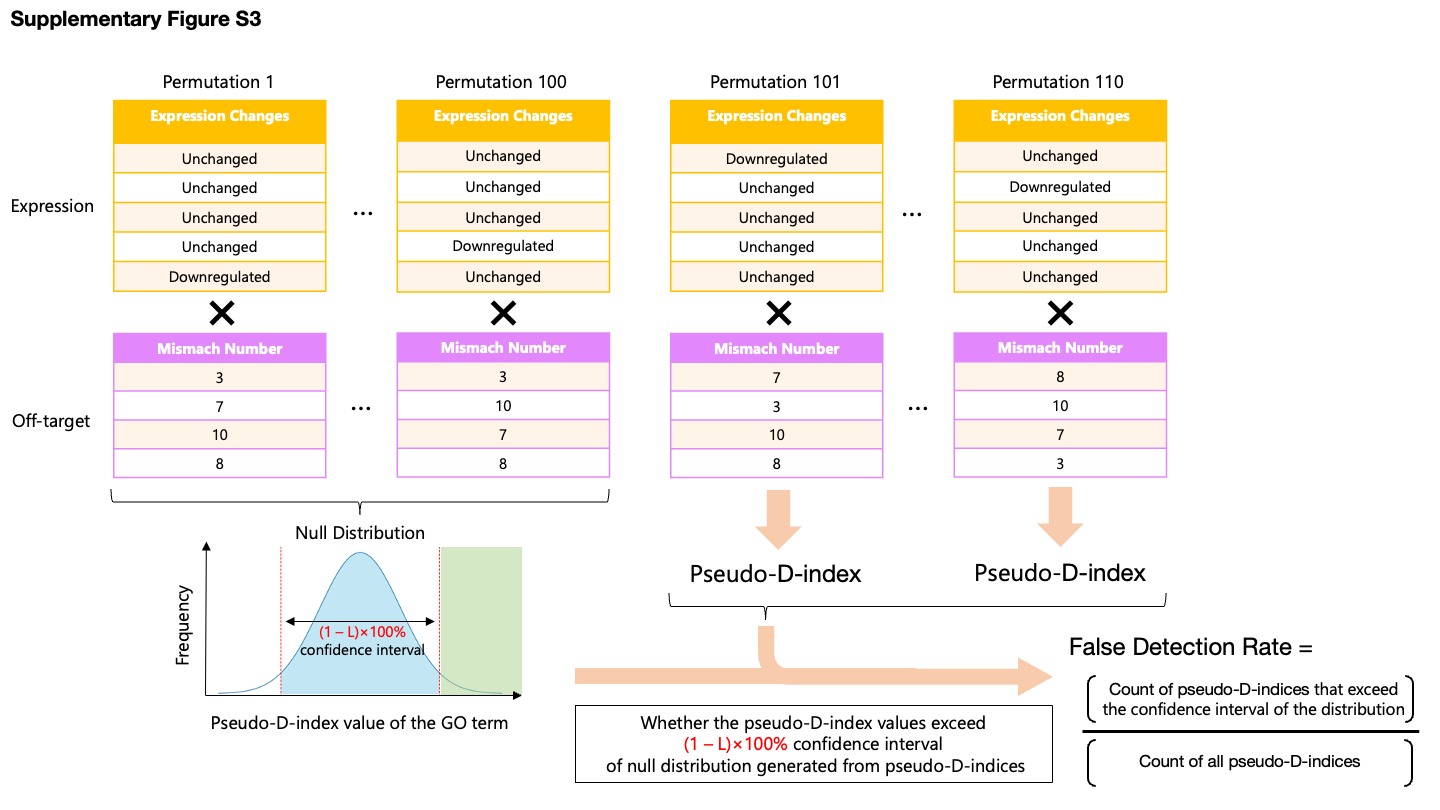

### Supplemental Figure 4

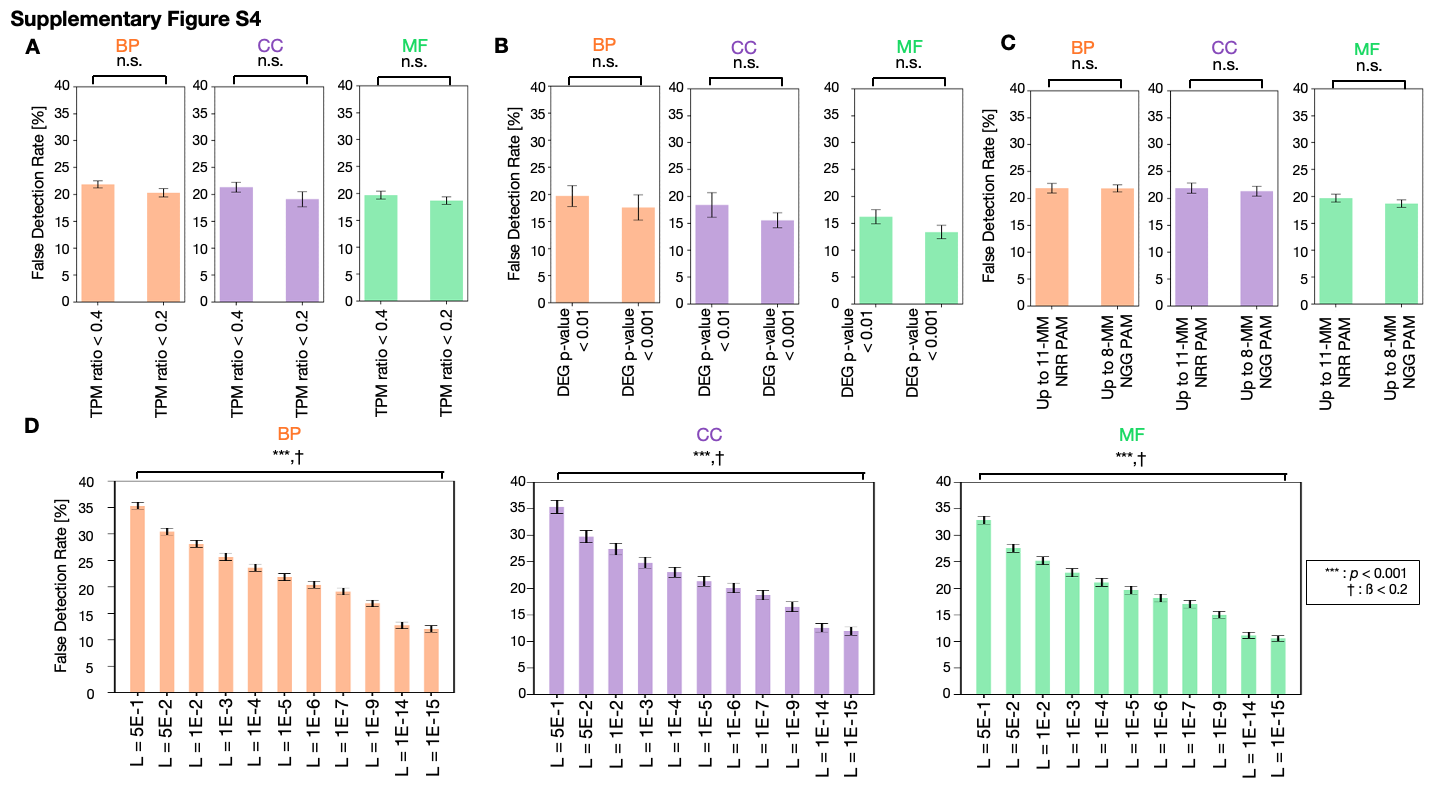
